# Molecular insights into ligand recognition and signaling of GPR99/OXGR1

**DOI:** 10.1101/2025.08.16.670657

**Authors:** Aijun Liu, Yezhou Liu, Yuanzhengyang Long, Richard D. Ye

## Abstract

GPR99/OXGR1 is a G protein-coupled receptor (GPCR) with two endogenous agonists, the tricarboxylic acid cycle derivative oxoglutarate and the inflammatory lipid mediator leukotriene E4 (LTE_4_). How GPR99/OXGR1 recognizes two distinct ligands is a biologically important question relevant to therapeutic development due to its established roles in asthma, allergy and inflammation. Here we present cryo-EM structures of GPR99/OXGR1-Gq complex with oxoglutarate and LTE_4_, respectively. The oxoglutarate-bound structure shows a binding pocket surrounded by the transmembrane domains (TM), with a primary site and an accessory site for simultaneous binding of two oxoglutarate molecules for full activation of the receptor. The TM binding pocket, however, is too small to accommodate the “Y” shaped structure of LTE_4_, a cysteinyl leukotriene. Alanine substitution of key residues for oxoglutarate binding had little impact on LTE_4_-induced signaling. An alternative site in between TM3/4/5 just above intracellular loop 2 was identified in the cryo-EM structure of GPR99/OXGR1-Gq complex formed with LTE_4_, but the densities were less well defined. Alanine substitution of amino acids potentially involved in LTE_4_ interaction at this site abrogated LTE_4_-induced receptor activation, with no effect on oxoglutarate-induced signaling. Both ligands activated GPR99/OXGR1 primarily through the Gq pathway, but LTE_4_ also induced inhibition of cAMP accumulation that was sensitive to pertussis toxin. These findings illustrate the structural basis for GPR99/OXGR1 interaction with oxoglutarate and suggest the presence of a distinct binding site for LTE_4_.

## INTRODUCTION

GPR99 was initially identified as a homologue of P2Y1, a purinergic GPCR that binds nucleotide ligands ^1^. Further studies found that GPR99 does not bind nucleotide ligands ^2^ but recognize endogenous dicarboxylates such as oxoglutarate ^3^. Oxoglutarate is mainly derived from the tricarboxylic acid (TCA) cycle and serves as an essential myometabokine for autocrine and paracrine signaling, as well as an immunometabolite ^4^. GPR99 recognizes oxoglutarate with an EC_50_ of approximately 200 μM and was subsequently named OXGR1 ^5^. GPR99/OXGR1 is expressed in various human tissues including the kidney, respiratory epithelium, placenta, muscle, fetal brain and immune cells ^1,4,6,7^. As a receptor of oxoglutarate, GPR99/OXGR1 is characteristic with its low affinity and low potency for this ligand, which is present in various tissues at concentrations in the micromolar range. In renal tubules, high oxoglutarate concentrations promote Gq-mediated IP1 accumulation, regulating acid-base homeostasis in kidney ^4^. Variations in renal pH may alter the polarity of oxoglutarate, thereby influencing Gq signaling through the activation of GPR99/OXGR1 ^4,8^. These context-dependent features highlight how physiological environments may shape GPR99/OXGR1 signaling.

In addition to oxoglutarate, leukotriene E_4_ (LTE_4_) was identified as a ligand of GPR99/OXGR1 ^9^. LTE_4_ is a cysteinyl leukotriene with a branched structure and established functions in allergy and asthma ^10,11^. Elevated concentrations of LTE_4_ are observed in severe asthma, acute exacerbation of asthma, and aspirin-exacerbated respiratory disease (AERD) ^12–14^, where it mediates sustained tracheobronchial constriction. An enrichment of GPR99/OXGR1 expression on the respiratory epithelial cells further exacerbates the inflammatory condition. An early study on LTE_4_-induced bronchoconstriction suggested that the high stability of LTE_4_, compared to LTC_4_ and LTD_4_ that are cysteinyl leukotrienes binding to different receptors, contributes to asthma development due to the chronic effect of receptor/ligand-related hyperresponsiveness ^15^. There is also experimental evidence pointing to the presence of the third CysLT receptor ^16^. Thus, GPR99/OXGR1 is recognized as the third identified cysteinyl leukotriene GPCR (CysLT_3_R), following CysLT_1_R and CysLT_2_R that primarily bind to LTC_4_ and LTD_4 5,9,17_. CysLT_3_R (GPR99/OXGR1) is markedly different from CysLT_1_R and CysLT_2_R in primary sequence with only 36% of amino acid identity ^9^. Moreover, the LTE_4_ concentrations required to activate CysLT_1_R and CysLT_2_R are much higher than its physiological levels, suggesting that the actions of LTE_4_ are primarily mediated through the third receptor ^17^. Furthermore, the binding affinity of LTE_4_ to GPR99 is higher than that of LTC_4_ and LTD_4_ to CysLT_1_R and CysLT_2_R, respectively ^7,9^. Extensive studies have associated LTE_4_-CysLT_3_R(GPR99/OXGR1) signaling with potent pro-inflammatory effects, such as enhanced bronchoconstriction ^3,18^ and eosinophil influx into the airway ^6,7,14^.

LTE_4_ is the second endogenous ligand of GPR99/OXGR1, with a high affinity for the receptor (K_d_ ∼ 2.5 nM) ^9^. This is in contrast to the low affinity of oxoglutarate (K_d_ in hundreds of micromolar), suggesting that the two structurally different ligands may interact with GPR99/OXGR1 in different manner. The structural basis for GPR99/OXGR1 to bind oxoglutarate and LTE_4_ is currently lacking, despite progress in structural characterization of CysLT_1_R and CysLT_2_R ^19–21^. Here we report a cryo-EM structure of GPR99/OXGR1 bound to oxoglutarate. Structural and functional analyses identify a primary and an accessory binding site for oxoglutarate, featuring simultaneous binding of two oxoglutarate molecules for full activation of the receptor. Furthermore, we found that the transmembrane binding pocket in GPR99/OXGR1 is too small to accommodate the branched LTE_4_, and further cryo-EM analysis of a GPR99/OXGR1-Gq complex formed in the presence of LTE_4_ identified a potential binding site outside the transmembrane binding pocket. Through functional analysis, we confirmed the alternative binding mode of LTE_4_ to GPR99/OXGR1, which likely serves as a promising target for future drug candidates. We found that LTE_4_ primarily activates the Gq-signaling pathways via GPR99/OXGR1, and the Gi pathway to a lesser extent. Although the poor EM density at the alternative site prevented us to definitively locate the LTE_4_ binding site, our study provides a rational framework of the structural basis for and mechanistic insights into ligand recognition of two different ligands by GPR99/OXGR1.

## RESULTS

### Overall structure of the oxoglutarate-GPR99/OXGR1-Gq signaling complex

The human GPR99/OXGR1 and Gq proteins were co-expressed in Sf9 insect cells and purified. The endogenous ligand oxoglutarate was introduced to facilitate the formation of the GPR99/OXGR1-Gq signaling complex, which was purified and subjected to cryo-EM analysis (Supplementary Fig. S1a,b). The structure of the signaling complex was then determined at a global resolution of 3.16 Å (Supplementary Fig. S1c-f, Table 1). The structure models were confidently built, and most residues were assigned based on the well-defined density map (Fig. 1a and Supplementary Fig. S2). Two separate ligand densities were observed in the orthosteric binding pocket, which fit well with two molecules of oxoglutarate (Fig. 1b). GPR99/OXGR1 holds the canonical seven-transmembrane architecture and G protein coupling manner of GPCR (Fig. 1c). There are two disulfide bridges in the extracellular domains: C24^N-term^-C274^7.25^, and C106^3.25^-C183^ECL2^ (using GPCR Ballesteros-Weinstein numbering as superscripts ^22^). The disulfide bonds are conserved among most Class A GPCRs and contribute to stabilization of the extracellular domains (Supplementary Fig. S3). The GPR99/OXGR1 in our structure adopts a conformation in the active state, with an intracellular pocket accommodating the α5 helix in Gq (Fig. 1b).

**Fig. 1.**
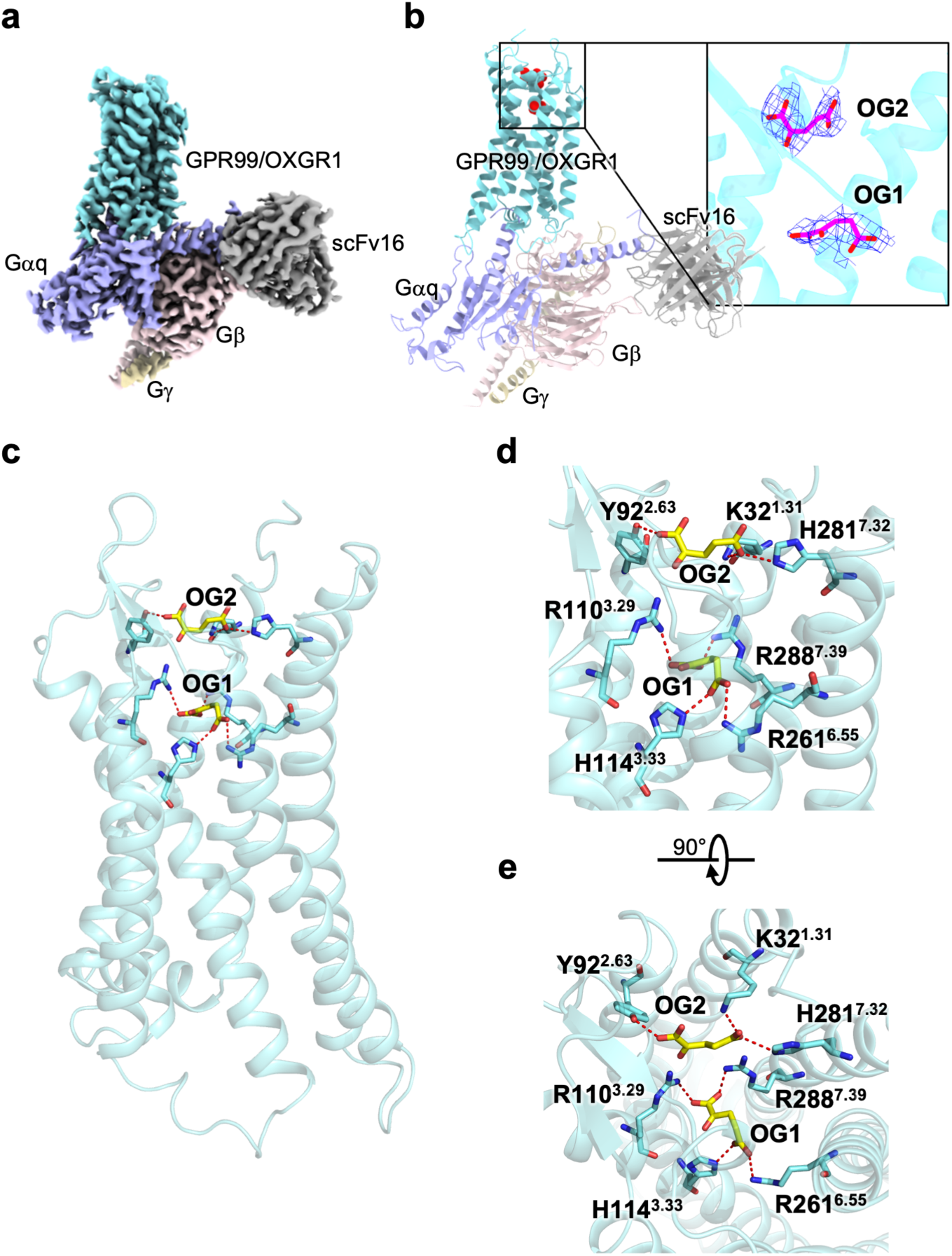
Overall structure of the oxoglutarate-GPR99/OXGR1-Gq complex. **a,** Cryo-EM density map of the oxoglutarate-GPR99/OXGR1-Gq complex. GPR99/OXGR1 is shown in cyan, Gαq in light purple, Gý in light pink, Gψ in wheat yellow, and scFv16 in grey. **b,** 3-D structure of oxoglutarate-GPR99/OXGR1-Gq complex (side view). The structure and EM density of the two oxoglutarate molecules (OG1, OG2) are highlighted in the inset. **c,** An enlarged view of OG1 and OG2 in the receptor, shown as yellow sticks. **d-e,** Molecular interactions between OG1 and OG2 and GPR99/OXGR1, in side view (**d**) and extracellular view (**e**). Polar interactions with key residues of the receptor are shown as red dashed lines.

**Table 1.** Cryo-EM data collection, model refinement and validation statistics.

| **Data collection and processing** | **Oxoglutarate-**<br>**GPR99/OXGR1-Gq** | **LTE<sub>4</sub>-**<br>**GPR99/OXGR1-Gq** |
| --- | --- | --- |
| Magnification | 105,000 | 105,000 |
| Voltage (kV) | 300 | 300 |
| Electron exposure (e<sup>-</sup>/Å<sup>2</sup>) | 52.0 | 52.0 |
| Defocus range (μm) | -1.2~-2.5 | -1.2~-2.5 |
| Pixel size (Å) | 0.85 | 0.85 |
| Symmetry imposed | C1 | C1 |
| Initial particle projections (no.) | 2,676,089 | 2,043,778 |
| Final particle projections (no.) | 167,941 | 100,083 |
| **Refinement** |  |  |
| Initial model used (Alpha Fold database code) | AF-Q96P68-F1 | AF-Q96P68-F1 |
| Map resolution (Å) | 3.16 | 2.84 |
| FSC threshold | 0.143 | 0.143 |
| **Model composition** |  |  |
| Non-hydrogen atoms | 9,097 | 9,103 |
| Protein residues | 1,153 | 1,152 |
| **R.m.s. deviations** |  |  |
| Bond lengths (Å) | 0.004 | 0.004 |
| Bond angles (°) | 0.973 | 0.596 |
| **Validation** |  |  |
| MolProbity score | 1.69 | 1.60 |
| Clashes core | 6.85 | 3.70 |
| **Ramachandran plot** |  |  |
| Favored (%) | 95.52 | 96.66 |
| Allowed (%) | 4.48 | 3.34 |
| Outliers(%) | 0.00 | 0.00 |

### Molecular basis for oxoglutarate recognition by GPR99/OXGR1

In the oxoglutarate-GPR99/OXGR1-Gq complex, the transmembrane domains TM1-3, TM6-7, extracellular loop (ECL) 1 and ECL2 of GPR99/OXGR1 encircle a hydrophilic and positively charged orthosteric pocket suited for the binding of negatively charged oxoglutarate molecule (Supplementary Fig. S4). The oxoglutarates adopt a horizontal orientation (Supplementary Fig. S4). One oxoglutarate molecule (OG1 in Fig. 1b) is embedded at the bottom of the pocket with the carboxyl group pointing downwards, and the other oxoglutarate molecule (OG2) fits well in the solvent-accessible subpocket at the top of the TM pocket with the carboxyl group pointing upwards (Fig. 1b and Supplementary Fig. S4). OG1 is directly clamped by salt bridges with four positively charged amino acids: R110^3.29^ and R288^7.39^ for one carboxyl, and H114^3.33^ and R261^6.55^ for the other (Fig. 1d,1e). There are sufficient interactions between OG1 and these amino acid residues that form a primary binding site for the ligand. In comparison, OG2 is salt-bridged with K32^1.31^ and H281^7.32^ at one carboxyl, and a hydrogen bond with Y92^2.63^ through the other carboxyl (Fig. 1d, 1e). The surrounding amino acids form an accessory site near ECL2, which is well exposed to the solvent environment and therefore less stable for OG2 binding. Notably, two gating arginine residues (R110^3.29^ and R288^7.39^), together with Y93^2.64^ and D185^ECL2^, spatially separate OG1 from OG2 (Supplementary Fig. S5a). The interaction network of arginines ^3^ stabilizes the extracellular domain of GPR99/OXGR1 and separates OG1 from the extracellular solvent environment (Supplementary Fig. S5b).

### Structural basis of selectivity for GPCR binding of dicarboxylates

Our findings reveal a distinctive binding mode for dicarboxylates in GPR99/OXGR1 compared to GPR91, suggesting a molecular mechanism underlying dicarboxylate selectivity in these GPCRs. The essential residues K32^1.31^, H281^7.32^ and Y92^2.63^ for oxoglutarate binding in GPR99/OXGR1 are not found in GPR91 (E22, N274 and S82 in GPR91). (Fig. 2 and Supplementary Fig. S6). Positively charged arginine residues are critical for the recognition and binding of negatively charged carboxyl groups, which are present in oxoglutarate (from oxoglutaric acid) and succinate (from succinic acid) in physiological pH. Most arginine residues in the orthosteric pocket are conserved between GPR91 and GPR99/OXGR1, namely R110^3.29^, R261^6.55^, R264^6.58^, R268^ECL3^, and R288^7.39^, but R177^ECL2^ and R180^ECL2^ in GPR99/OXGR1 are absent in GPR91 (Fig. 2 and Supplementary Fig. S6). In GPR91, only R99^3.29^ and R281^7.39^ are known to participate in direct interactions with identified agonists and antagonists, including succinate, epoxysuccinate, maleic acid, compound 31, and NF-56-EJ40 ^23,24^.

**Fig. 2.**
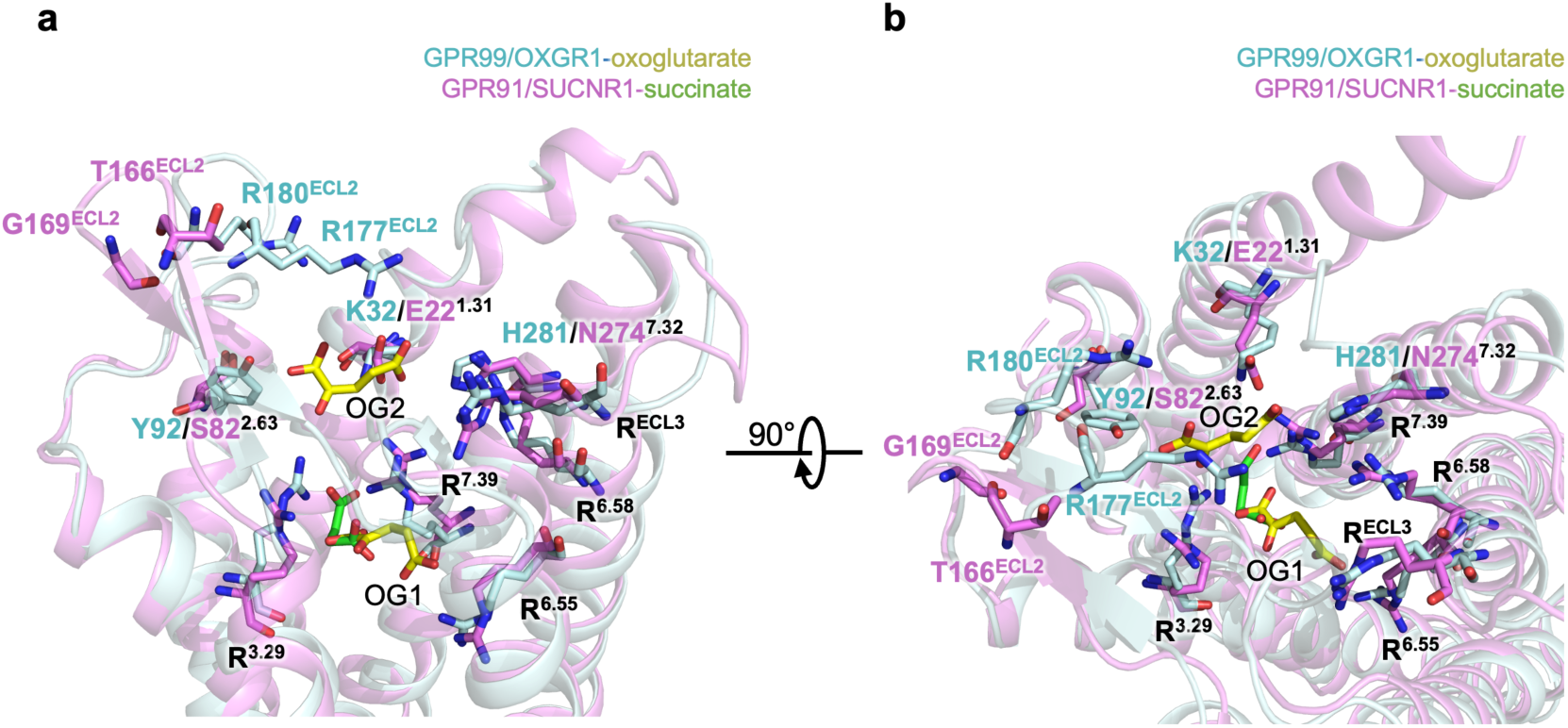
Comparison of the dicarboxylate ligand binding pockets of GPR99/OXGR1-oxoglutarate and GPR91/SUCNR1-succinate. Side view (**a**) and top view (**b**) of the transmembrane binding pockets of GPR99/OXGR1 (cyan) bound with oxoglutarate (yellow) and GPR91/SUCNR1 (pink) bound with succinate (green). Ligands and key residues for ligand binding are shown as sticks.

We found that, in GPR99/OXGR1, R110^3.29^, R261^6.55^ and R288^7.39^ contributed to the binding of OG1 and were involved in receptor activation. The other four arginine residues, R177^ECL2^, R180^ECL2^, R264^6.58^ and R268^ECL3^, were positioned above the OG1 and at the top of the orthosteric pocket, with their positively charged guanidinium groups oriented down towards the orthosteric pocket (Supplementary Fig. S5b). This cluster of arginines likely acts as a gatekeeper of the orthosteric pocket, facilitating the recruitment of oxoglutarate. In particular, the flexibility of R177^ECL2^ and R180^ECL2^ at the tip of ECL2 in GPR99/OXGR1 may be crucial for capturing negatively charged molecules, allowing only appropriate ligands like oxoglutarate to engage with R264^6.58^ and R268^ECL3^ (Supplementary Fig. S5b). Interestingly, the positively charged R177^ECL2^ and R180^ECL2^ in GPR99/OXGR1 are replaced by the negatively charged D167^ECL2^ and N168^ECL2^ in GPR91. Additionally, the tip of ECL2 in GPR91 is positioned farther from the orthosteric pocket compared to GPR99/OXGR1. These observations suggest differences in ligand recognition and recruitment between GPR99/OXGR1 and GPR91, thus contributing to ligand selectivity in these two dicarboxylate receptors. The closely positioned arginine residues at the orifice and surface of the orthosteric binding pocket form an “arginine network” that guides and mediates the recruitment and binding of oxoglutarate to the receptor.

### Structural insights into LTE_4_ binding to GPR99/OXGR1

Unlike oxoglutarate, LTE_4_ has a relatively large, hydrophobic tail and two charged carboxyl groups at the head (Fig. 3a). Like other CysLTs, the “Y” shaped carboxyl terminus of LTE_4_ differs from the straight chain of hydroxyacids found in LTB_4 10,25_. To learn how LTE_4_ binds GPR99/OXGR1, we compared our solved structure with available structures of CysLT_1_R and CysLT_2_R ^19,20^, bound to Pranlukast (Fig. 3b) and ONO-2570366 (Fig. 3c), respectively. Amino acid residues known to interact with these ligands were identified with the superimposed receptor structures shown in the background. The binding pockets showed little resemblance at these critical residues, an observation consistent with the grouping of GPR99/OXGR1 in the P2Y nucleotide receptor subfamily and the two CysLT receptors in a different subfamily ^1,26^. We also compared the receptor structures sliced at the TM binding pocket in the same plane. As shown in Fig. 3d-3f, the binding pockets of CysLT_1_R and CysLT_2_R are similar in shape and volume (Fig. 3d, 3e), snugly holding their respective ligands. In comparison, the binding pocket of GPR99/OXGR1 differs drastically from these two receptors in shape and volume, suitable only for the much smaller oxoglutarate molecules. Other notable features include the absence of positively charged arginine network, which is important for binding of the bicarboxylic ligands such as oxoglutarate and succinate ^3,24,27^, from the CysLT receptors. Moreover, the large access cleft from the lipid membrane between TM4 and TM5 in the CysLT receptors ^21,26^, which allow lateral entry of the bulky tails of the lipid ligands, is missing from GPR99/OXGR1. While revising the manuscript, we compared the structure of GPR99/OXGR1 with the LTD_4_-bound CysLT2R structure that became available only recently ^21^, and found little resemblance between the two at the orthosteric binding pockets (Supplemental Fig. S7). These results strongly suggest that LTE_4_ uses a different binding site in GPR99/OXGR1.

**Fig. 3.**
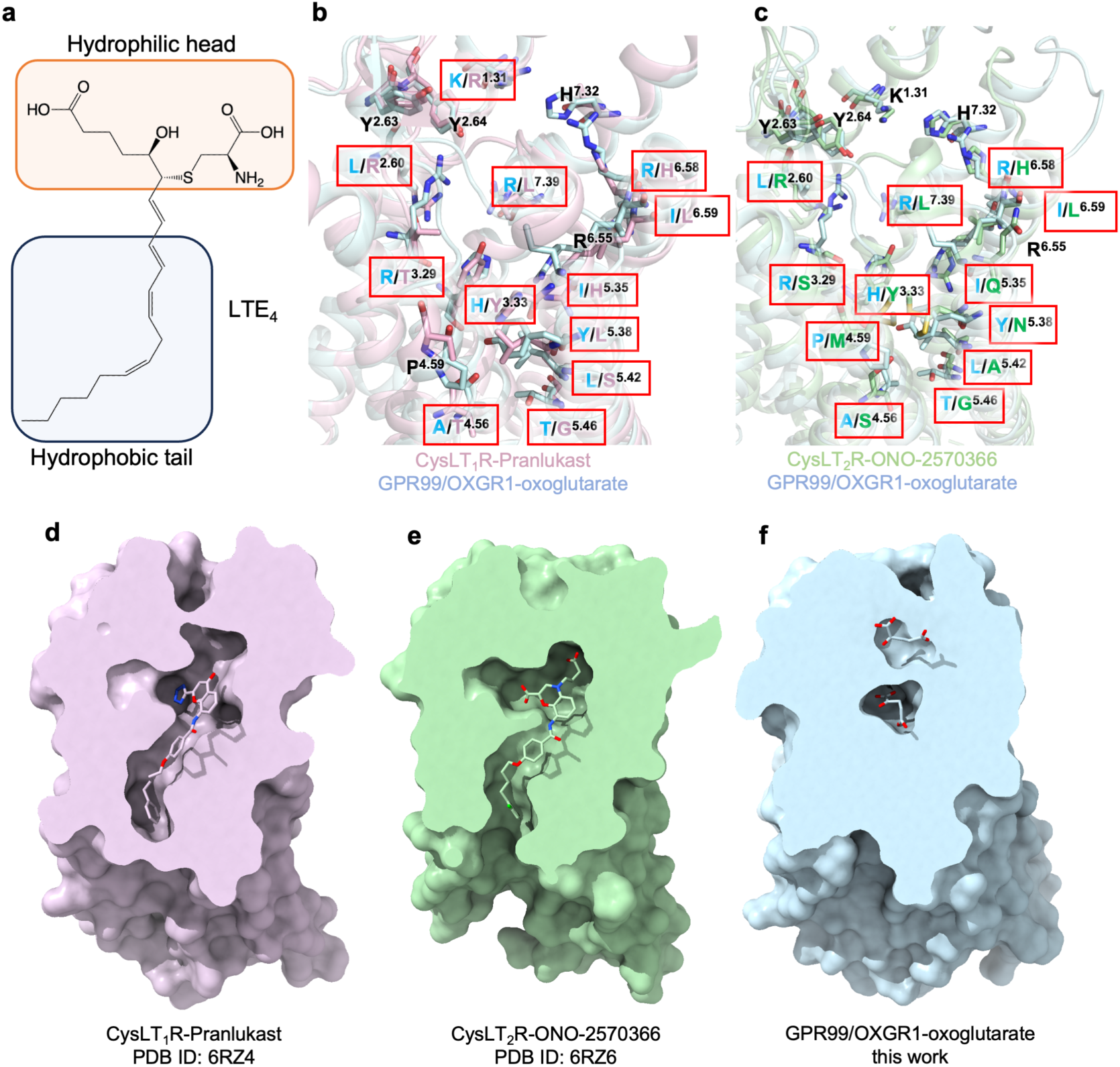
Comparison of ligand binding pockets of CysLT_1_R-Pranlukast, CysLT_2_R-ONO-2570366 and GPR99/OXGR1-oxoglutarate structures. **a,** Chemical structure of the LTE_4_ molecule, the hydrophilic head and the hydrophobic tail of LTE_4_ are highlighted in boxes. **b,** Comparison of TM binding pocket residues in superimposed structures of CysLT_1_R (light pink, PDB ID: 6RZ4) and GPR99/OXGR1 (light cyan, this work). **c,** Comparison of TM binding pocket residues in superimposed structures of CysLT_2_R (light green, PDB ID: 6RZ6) and GPR99/OXGR1 (light cyan, this work). **d-f,** Superimposed receptor structures of CysLT_1_R-Pranlukast, CysLT_2_R-ONO-2570366 and GPR99/OXGR1-oxoglutarate, sliced at the TM binding pocket in the same plane. GPR99/OXGR1-oxoglutarate structure is aligned as a reference model for structural comparison.

To illustrate how LTE_4_ binds to GPR99/OXGR1, an LTE_4_-GPR99/OXGR1-Gq complex was obtained with an excessive amount of LTE_4_ (∼1.00 μM), and then subjected to cryo-EM analysis (Supplementary Fig. S8a,b). The resulting structure of the complex was resolved at an overall resolution of 2.84 Å (Fig. 4a, Supplementary Fig. S8c-g and S9, Table 1). An analysis of the structure found no extra EM density for LTE_4_ in the orthosteric pocket. Instead, an alternative binding site was identified above the intracellular loop 2 (ICL2) and surrounded by TM3-5. Although the EM density could not definitively indicate binding of LTE_4_ at this site, the shape fits well with an LTE_4_ molecule (Fig. 4b,c).

**Fig. 4.**
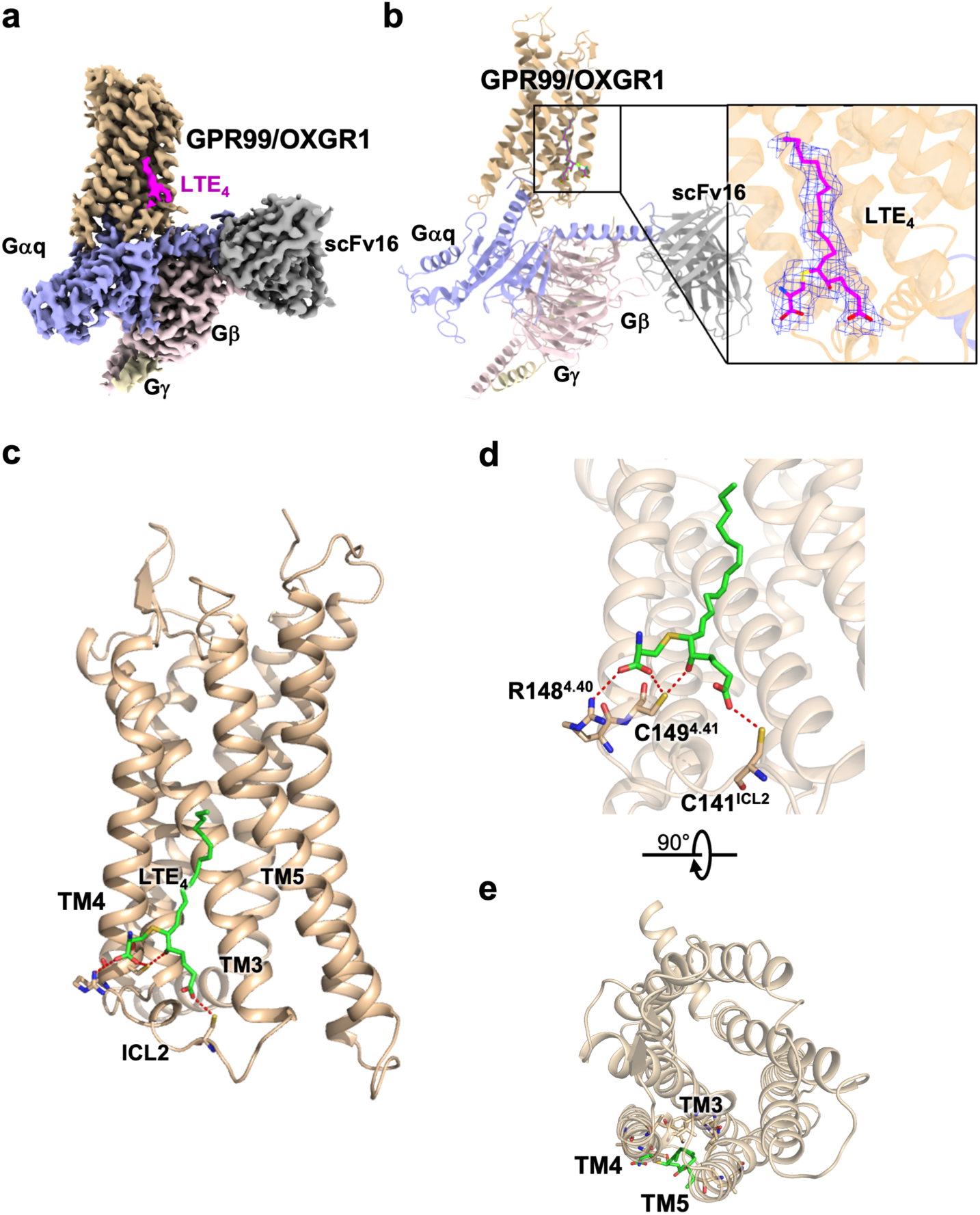
Overall structure of the LTE_4_-GPR99/OXGR1-Gq complex. **a,** Cryo-EM density map of the LTE_4_-GPR99/OXGR1-Gq complex. GPR99/OXGR1 is shown in wheat, Gαq in light purple, Gý in light pink, Gψ in wheat yellow, and scFv16 in grey. **b,** 3-D structure of the LTE_4_-GPR99/OXGR1-Gq complex (side view). The structure and EM density (inset) of the bound LTE_4_ are highlighted on the right. **c,** An enlarged view of the LTE_4_ bound to the receptor. **d-e,** Molecular interactions between LTE_4_ (green) and GPR99/OXGR1, in side view (**d**) and extracellular view (**e**). Polar interactions with key residues of the receptor are shown as red dashed lines.

Several approaches were taken to further explore the alternative site for LTE_4_. A structural complex model was built based on the EM density, and potential interactions between LTE_4_ and GPR99/OXGR1 were identified. Specifically, L122^3.41^, T125^3.44^, I129^3.48^, F130^3.49^, V152^4.44^, A153^4.45^, V157^4.49^, I160^4.52^, T205^5.45^, L209^5.49^ could form extensive hydrophobic interactions with the aliphatic chain of LTE_4_ (Supplementary Fig. S10a). The negatively charged head of LTE_4_ could be attached to a positively charged subpocket formed by ICL2 and TM3-4 (Fig. 4d and Supplementary Fig. S10b). In detail, R148^4.40^ could form a salt bridge with one carboxyl group, C141^ICL2^ could possibly form a hydrogen bond with the other carboxyl group, and C149^4.41^ could form a hydrogen bond with the hydroxyl group as well as another hydrogen bond with a carboxyl group (Fig. 4d, 4e). The presence of these interactions was validated with alanine substitutions and the effects on receptor signaling by LTE_4_ as well as oxoglutarate were compared (see below).

Computational approaches were taken to further evaluate possible binding of LTE_4_ to this alternative site. Molecular docking was applied to dock LTE_4_ to the TM pocket and the alternative site, respectively. The corresponding binding energies were calculated, showing an energetic preference for LTE_4_ at the alternative binding site (Supplementary Fig. S11a). Furthermore, three independent 1-µs MD simulations captured a stable binding pose of LTE_4_ at this alternative site (Supplementary Fig. S11b-c).

### Validation of GPR99/OXGR1-ligand interactions by site-directed mutagenesis

To further elucidate ligand interaction with GPR99/OXGR1, site-directed mutagenesis and functional assays were performed. Amino acid residues exhibiting polar interactions with oxoglutarate and LTE_4_ were substituted by alanine, and all mutants were positive for cell surface expression (Supplementary Fig. S12a). The accumulation of IP1, downstream of the Gq-phospholipase C pathway, was measured for both WT and mutants of GPR99/OXGR1 (Fig. 5). Alanine substitution of residues interacting with OG1 (R110A, H114A, R261A, R288A) resulted in a significant reduction in both the potency and efficacy of IP1 accumulation following oxoglutarate stimulation (Fig. 5a). In comparison, mutations in OG2-interacting residues (K32A and Y92A) led to a more than 100-fold increase in EC_50_, while the maximum response (E_max_) remained unchanged (Fig. 5a). These findings suggest that while OG2 is essential for maintaining ligand occupation, it plays a less significant role in receptor activation compared to OG1.

**Fig. 5.**
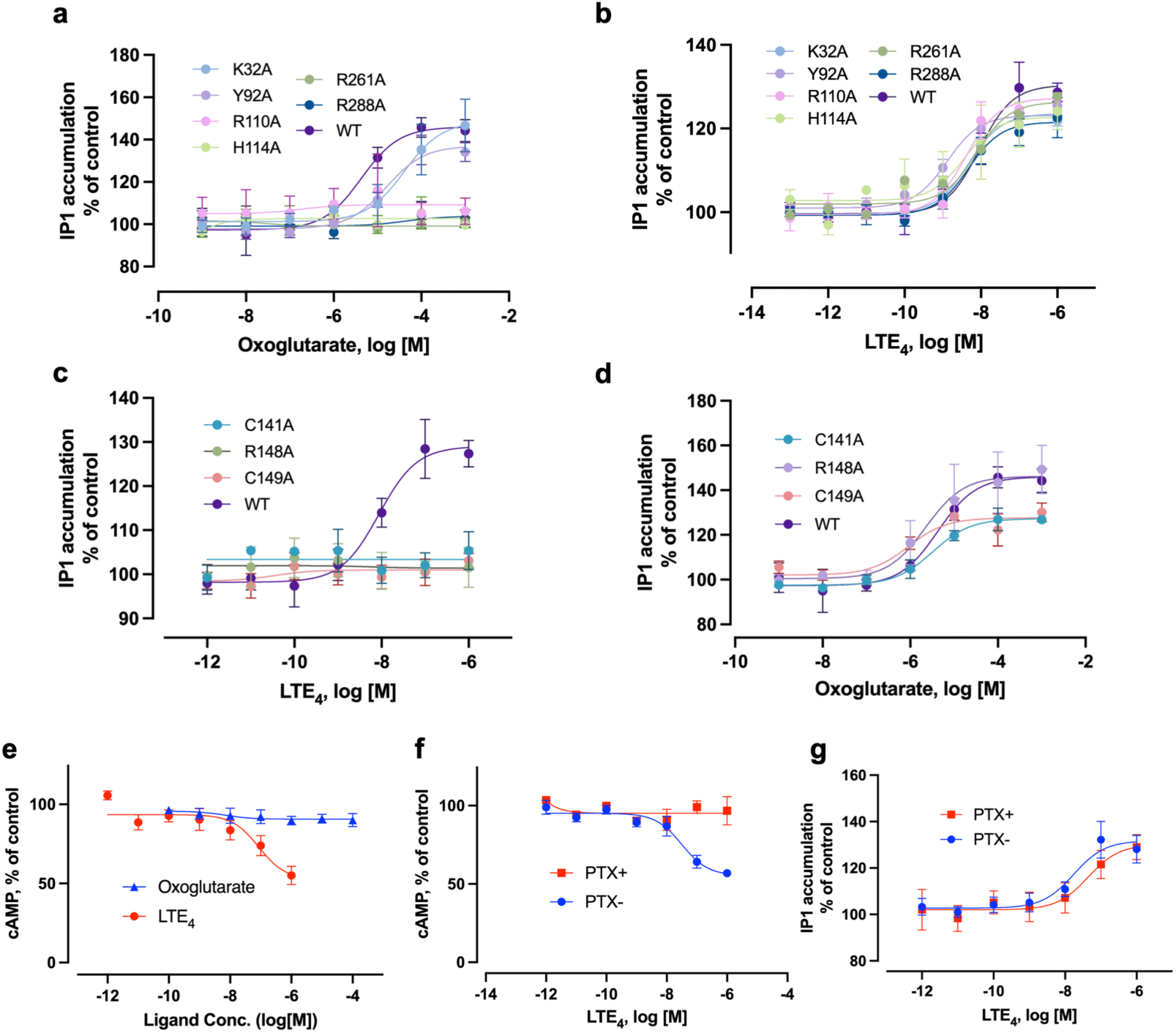
Comparison of WT and mutant GPR99/OXGR1 receptors in functional assays. Transfected cells (HeLa) expressing either WT or mutant receptors were stimulated with oxoglutarate or LTE_4_, the induced responses were measured to reflect G protein activation (cAMP reduction for Gαi, IP1 accumulation for Gαq). **a,** 6 mutants with alanine substitutions of key residues for oxoglutarate binding were tested against WT in IP1 accumulation. Four of the 6 mutants abrogated GPR99/OXGR1 signaling for Gαq activation (IP1 accumulation), and the other 2 mutants reduced the response. **b,** the same mutants had no or little effect on LTE_4_-induced IP1 accumulation. **c** and **d,** 3 alanine-substituted mutants of residues suspected to interact with LTE_4_ at the projected site were tested against WT in IP1 accumulation assay. All 3 mutants abrogated the LTE_4_-induced IP1 accumulation (**c**) while having little or much smaller effect on oxoglutarate-induced response (**d**). In **e**, LTE_4_, but not oxoglutarate, stimulated cells expressing the WT GPR99/OXGR1 with reduction of cAMP, indicating Gαi activation. In **f**, pertussis toxin (PTX), a bacterial toxin that blocks Gαi activation, abrogated LTE_4_-induced cAMP reduction, supporting coupling of GPR99/OXGR1 with Gi proteins in addition to Gq. In **g**, PTX had no effect on LTE_4_-induced IP1 accumulation that does not exclusively rely on Gi in the presence of Gq proteins. Data are shown as mean ± SEM from three independent experiments.

Consistent with the predicted structure for alternative binding of LTE_4_, alanine substitution of the OG1 and OG2 sites had little effect on LTE4-induced receptor signaling (Fig. 5b). Substitution of the projected LTE_4_-binding residues (C141^ICL2^, R148^4.40^ and C149^4.41^) markedly reduced LTE_4_-induced IP1 accumulation (Fig. 5c), but had little effect on oxoglutarate-induced receptor activation (Fig. 5d). This finding suggests that LTE_4_ binding to GPR99/OXGR1 depends on hydrogen bonding with key residues C141^ICL2^, R148^4.40^ and C149^4.41^ in the alternative binding site, but this site was not utilized by oxoglutarate. Altogether, results obtained from mutagenesis experiments provide functional clues that support the binding of LTE_4_ to a site distinct from the orthosteric site for oxoglutarate. We have also noticed that some mutants (C141^ICL2^A, C149^4.41^A) exhibited a reduced E_max_ albeit having comparable cell surface expression with the WT GPR99/OXGR1. As C141^ICL2^ and C149^4.41^ are proximal to the ICL2 region where the receptor extensively interacts with G proteins, we suspect that mutants of these sites may alter the overall stability of ICL2 and consequently the recruitment and activation of G proteins, and affect the efficiency of Gq response. These functional data distinguish the orthosteric binding mode of oxoglutarate from the alternative binding mode of LTE_4_ at GPR99/OXGR1.

### Ligand-induced signaling through GPR99/OXGR1

Given the structural difference between oxoglutarate binding and LTE_4_ binding at GPR99/OXGR1, it is suspected that their downstream signaling properties might vary. We therefore examined the signaling pathways triggered by oxoglutarate and LTE_4_. Notably, oxoglutarate preferentially activated the PLC pathway downstream of Gq signaling, leading to increased IP1 accumulation; however, oxoglutarate failed to stimulate cAMP reduction downstream of Gi coupling (Fig. 5e). In contrast, LTE_4_ treatment caused reduction in cAMP accumulation (Fig. 5e), suggesting activation of Gαi. This interesting finding was further verified by treatment of the cells with pertussis toxin (PTX), which ADP-ribosylates the Gαi and breaks the Gi activation cycle ^28^. PTX abrogated the cAMP response elicited by LTE_4_ (Fig. 5f) without affecting IP1 accumulation (Fig. 5g). These findings indicate that LTE_4_ binding to GPR99/OXGR1 can engage Gi-mediated signaling in addition to its Gq-coupled activity.

To investigate GPR99/OXGR1 basal activity, we used a NanoBiT-based G protein dissociation assay, with SmBiT fused to Gγ and LgBiT fused to Gα ^29^. Gα dissociation from Gβγ, that accompanies G protein activation, reduces the NanoBiT signal. HEK293T cells were co-transfected with equal amounts of plasmids encoding tagged G proteins and varying amounts of plasmids encoding GPR99/OXGR1 or the constitutively active KSHV-GPCR as a control ^30^. Basal G protein dissociation of GPR99/OXGR1 was compared to that of KSHV-GPCR. Increasing KSHV-GPCR plasmid levels resulted in a dose-dependent decrease in NanoBiT signal, indicating agonist-independent G protein dissociation, whereas GPR99/OXGR1 showed no signal change, suggesting negligible basal activity of GPR99/OXGR1 (Supplementary Fig. S12b). These results indicate that GPR99/OXGR1 is not an auto-activating GPCR that can complex with G proteins without agonist binding.

### The G protein coupling interface of GPR99/OXGR1

The G protein interfaces are highly similar between oxoglutarate-bound and LTE_4_-bound GPR99/OXGR1 signaling complex. However, subtle differences are observed for sidechain orientations of the α5 helix of Gq protein (Fig. 6a). As a result, the G protein interaction profiles are slightly different between oxoglutarate-bound and LTE_4_-bound GPR99/OXGR1 structures. In the oxoglutarate-GPR99/OXGR1-Gq complex, more extensive polar interactions between Gq and GPR99/OXGR1 were identified (Fig. 6b). Specifically, Y351 of the α5 helix of Gq protein forms polar contacts with R131^3.50^ and S68^2.39^ of GPR99/OXGR1. E350 forms a hydrogen bond with H145^ICL2^, and K349 interacts with D307^8.48^. Q345 forms polar interactions with Q230^ICL3^, and D341 has a polar contact with K236^6.30^ (Fig. 6b). In the LTE_4_-GPR99/OXGR1-Gq complex, four pairs of polar contacts are found between GPR99/OXGR1 and the Gq protein: R131^3.50^-Y351, D307^8.48^-K349, V134^3.53^-N347 and K236^6.30^-D341 (Fig. 6c). These subtle structural differences may help to explain the distinct biological outcomes induced by oxoglutarate versus LTE_4_.

**Fig. 6.**
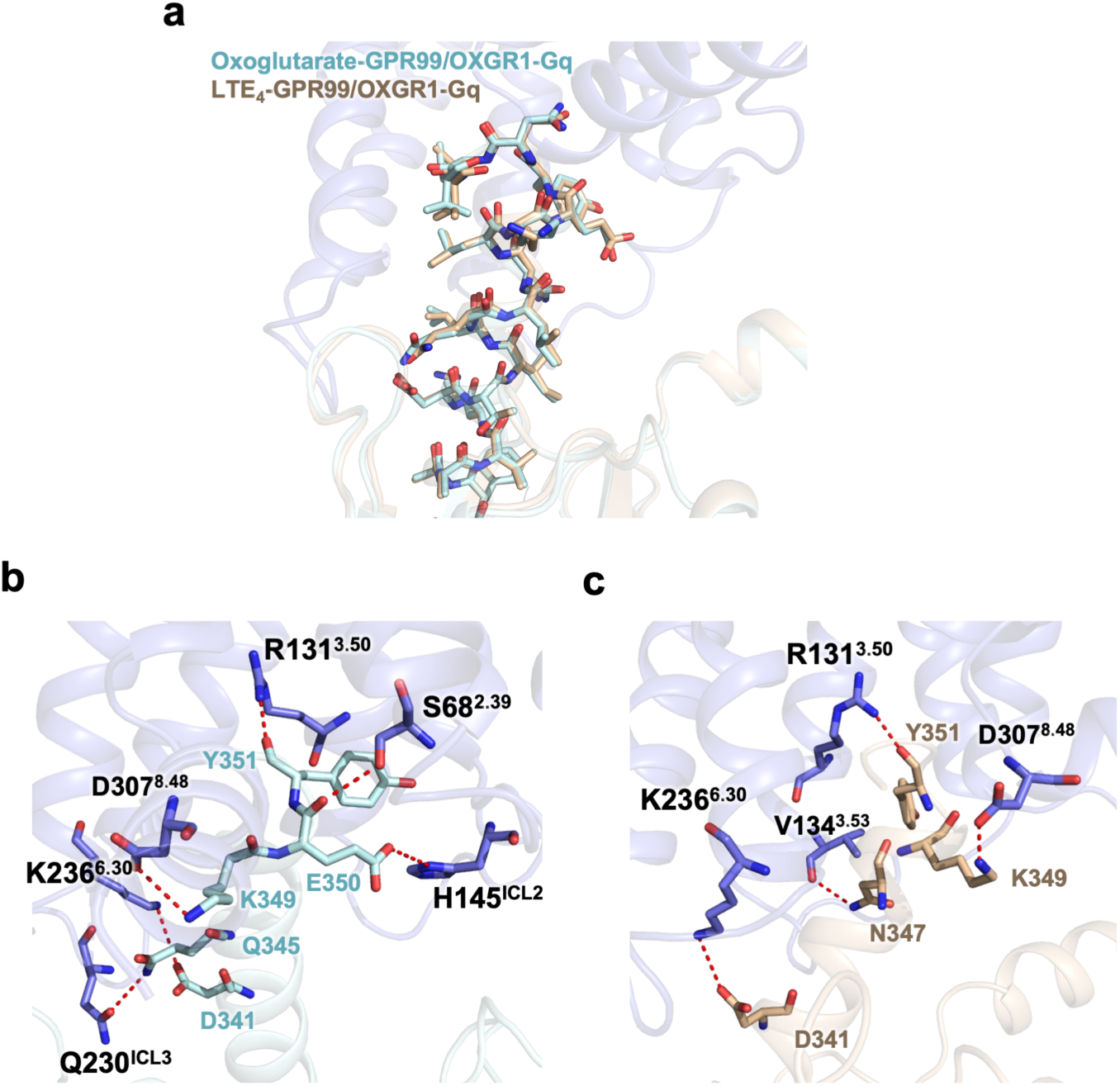
The G protein interface of oxoglutarate-bound and LTE_4_-bound GPR99/OXGR1. **a,** Superimposed structures of the Gq protein in the oxoglutarate-bound and LTE_4_-bound GPR99/OXGR1 signaling complex. Gq is shown in cyan for the oxoglutarate-bound structure and in champaign for the LTE_4_-bound structure. **b,** Molecular interactions between Gq and GPR99/OXGR1 in the oxoglutarate-bound complex. Oxoglutarate is shown in cyan, with polar interactions highlighted by red dashed lines. **c,** Molecular interactions between Gq and GPR99/OXGR1 in the LTE_4_-bound complex. LTE_4_ is shown in champaign, with polar interactions marked by red dashed lines.

### Ligand-induced activation mechanism of GPR99/OXGR1

To elucidate the activation mechanism induced by orthosteric coupling of oxoglutarate, the recently reported structures of GPR91 in the active state ^24^ (PDB: 8WOG) and inactive state ^23^ (PDB: 6RNK) were used as references for comparison. The outward movement of TM5-6 and inward movement of TM7, which accommodated the α5 helix of Gq, illustrated a typical conformational change associated with receptor activation (Fig. 7a,b). Additionally, the structural motifs responsible for GPCR activation, specifically DRY, CWXP, NPXXY and PIF (corresponding to FRY, CFLP, NLLLY and PIF in GPR99), were compared between GPR99/OXGR1 and GPR91 (PDB: 8WOG). The oxoglutarate-GPR99/OXGR1-Gq complex represented a molecular mechanism for receptor activation triggered from the orthosteric pocket. Specifically, the salt bridge between OG1 and R261^6.55^ gave rise to an inward turning of TM6, which was transduced through the CFLP motif (Fig. 7c). The turning of TM6, along with the bridge between OG1 and H114^3.33^, caused a rearrangement of the hydrophobic core of the PIF motif (Fig. 7d). Besides, the salt bridge between OG1 and R114^3.33^, and the hydrogen bond between OG1 and R288^7.39^, triggered a conformational change that propagates through TM7. The salt bridge between oxoglutarate and R110^3.29^ initiated conformational changes that were transduced through TM3. Furthermore, OG2 played an accessory role, alongside OG1, in initiating receptor signaling. In a synergistic manner, the extracellular signal induced a rearrangement of the NLLLY (NPXXY) motif (Fig. 7e) and the FRY(DRY) motif (Fig. 7f) in the intracellular domain, opening the intracellular pocket to accommodate α5 helix of the Gq protein (Fig. 7e, 7f).

**Fig. 7.**
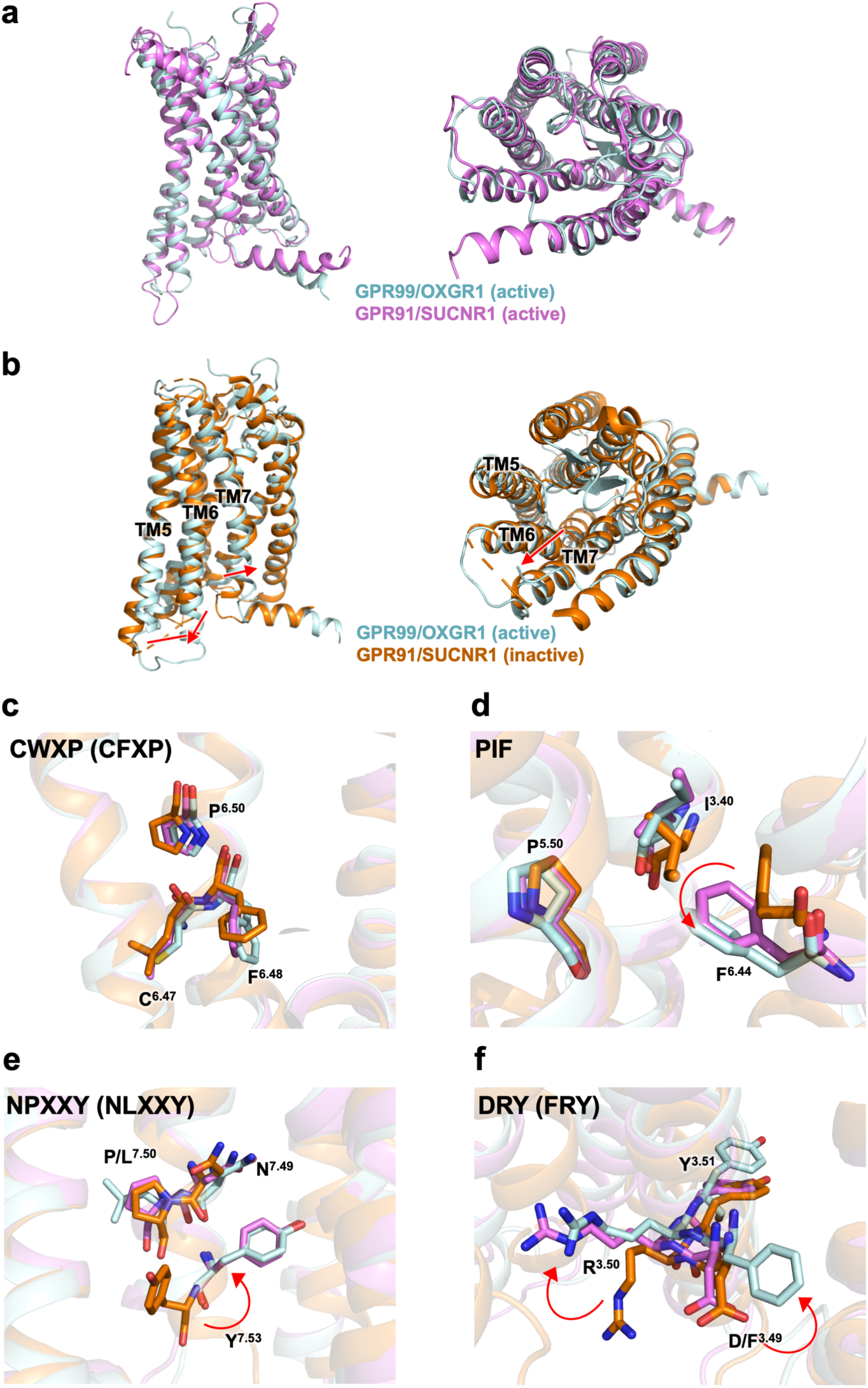
Mechanism of GPR99/OXGR1 activation induced by oxoglutarate binding to the orthosteric pocket. **a,** Superimposed structures of GPR99/OXGR1 (this study, oxoglutarate-bound) and GPR91/SUCNR1 in the active state (PDB ID: 8WOG). **b,** Superimposed structures of GPR99/OXGR1 (this study) and GPR91/SUCNR1 in the inactive state (PDB ID: 6RNK). **c-f,** Conformational changes at key activation motifs. Shown are side chain conformation at the CWxP (CFxP in GPR99/OXGR1 and GPR91/SUCNR1) motif (**c**), conformational changes at the PIF motif (**d**), conformational changes at the NPxxY (NLxxY in GPR99/OXGR1) motif (**e**), and rotameric conformation changes at the DRY (FRY in GPR99/OXGR1) motif (**f**). GPR99/OXGR1 in the active state is shown in cyan, GPR91/SUCNR1 in the active state is shown in pink, and GPR91/SUCNR1 in the inactive state is shown in orange.

The reported inactive state structures of CysLT_1_R and CysLT_2_R were used as references to illustrate the activation mechanism of GPR99/OXGR1 induced by LTE_4_ ^19,20^ (Fig. 8). In this structure, polar interactions caused the ICL2 of GPR99/OXGR1 to move outward when LTE_4_ is present (Fig. 8a), which induced the opening of the G protein-coupling pocket (Fig. 8b). The hydrophobic interaction driven by the aliphatic chain of LTE_4_ directly triggered the rearrangement of the NLxxY motif (Fig. 8c) and the FRY motif (Fig. 8d). Consequently, the CFxP and PIF motifs rearranged synergistically, with TM5-6 moving outward and TM7 moving inward (Fig. 8e,f). Additionally, a salt bridge formed between R318 and D312, further stabilizing the activated conformation and resulting in an ordered helix8, which was absent in the inactive structure due to its flexibility.

**Fig. 8.**
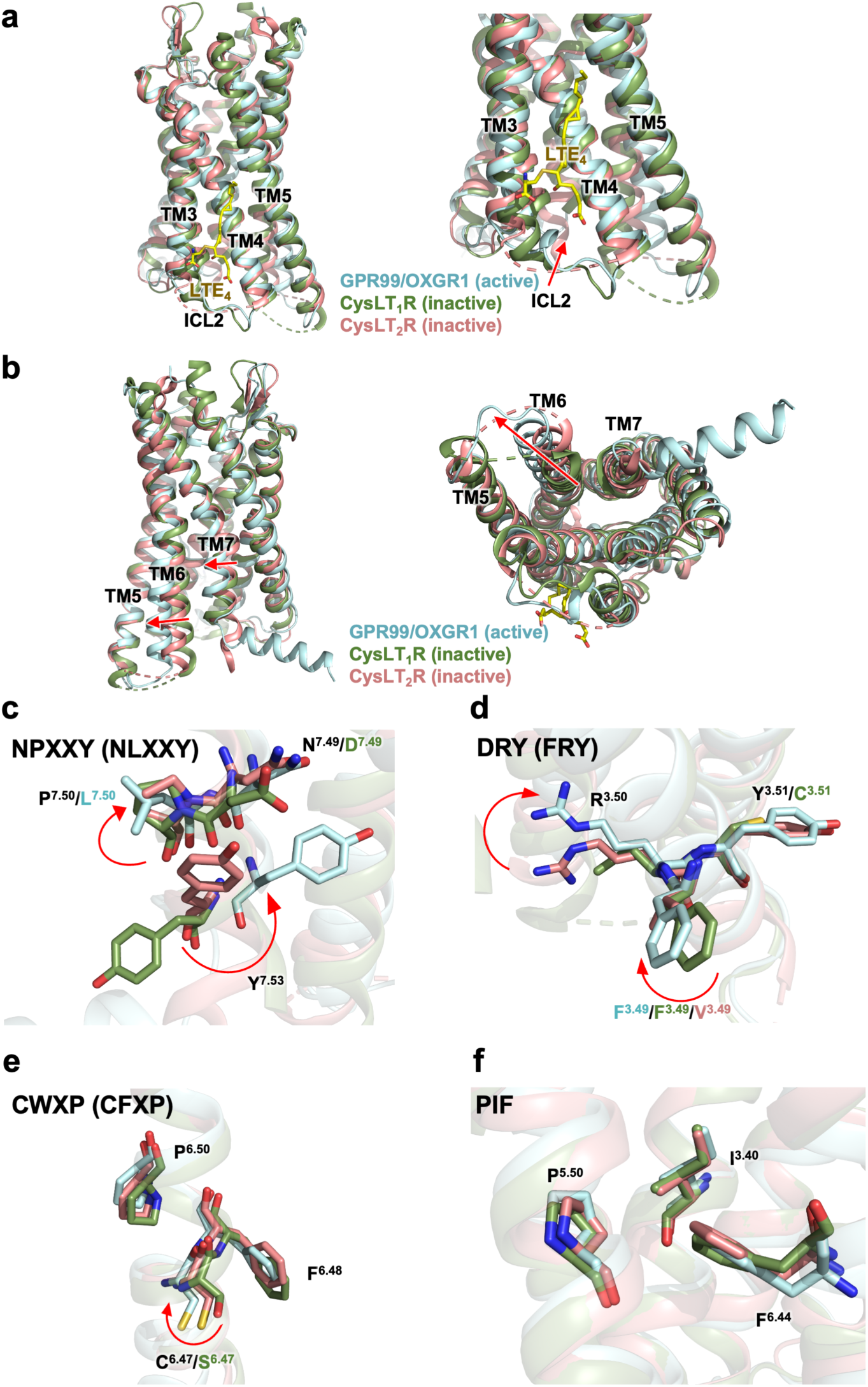
Mechanism of GPR99/OXGR1 activation induced by LTE_4_ binding to the non-canonical site. **a,** Superimposed structures of GPR99/OXGR1 (this study, LTE_4-_bound), CysLT_1_R in the inactive state (PDB ID: 6RZ4) and CysLT_2_R in the inactive state (PDB ID: 6RZ6). LTE_4_ is shown as yellow sticks. **b,** Superimposed structures of GPR99/OXGR1, inactive CysLT_1_R and inactive CysLT_2_R. Conformational changes are marked with red arrows. **c-f,** Conformational changes at key activation motifs. Shown are conformational changes at the NPxxY (NLxxY in GPR99/OXGR1, DPxxY in CysLT_1_R) motif (**c**), rotameric conformation changes at the DRY (FRY in GPR99/OXGR1, FRC in CysLT_1_R, VRY in CysLT_2_R) motif (**d**), conformational changes at the CWxP (CFxP in GPR99/OXGR1 and CysLT_2_R, SFxP in CysLT_1_R) motif (**e**), and conformation changes at the PIF motif (**f**). GPR99/OXGR1 (CysLT_3_R) in the active state is shown in cyan, CysLT_1_R in the inactive state is shown in green, and CysLT_2_R in the inactive state is shown in salmon pink.

## DISCUSSION

The structural mechanisms by which both dicarboxylates and CysLTs bind their native receptors remain underexplored. Persistent questions revolve around how GPCRs recognize small, charged dicarboxylates with high specificity but low efficacy. There have been great computational efforts, but the issues remain due to the lack of effective structural references ^31,32^. For example, an exhaustive metadynamics analysis identified two low-energy succinate binding sites in the orthosteric pocket of GPR91 ^31^. However, subsequent structural studies demonstrated the presence of only a single succinate binding site in this pocket ^24,27^. Furthermore, in the absence of a resolved structure for the oxoglutarate receptor GPR99/OXGR1, it remains challenging to address how dicarboxylates are selectively recognized by their respective receptors. This understanding is crucial for developing targeted drugs with high specificity.

In this study, we present a high-resolution cryo-EM structure of GPR99/OXGR1 bound to the dicarboxylate oxoglutarate. Our structural and functional analyses reveal the molecular basis for ligand selectivity and G protein coupling preferences. Dicarboxylates are critical components in physiological fluids, and as a result, their native GPCRs may have evolved orthosteric adaptations that attenuate or desensitize activation to prevent unnecessary receptor activation under normal conditions. Notably, GPR99/OXGR1 and GPR91/SUCNR1 have shared features including the arginine network at their binding pockets, but also distinct features in GPR99/OXGR1 allowing simultaneous binding of two oxoglutarate molecules. Full activation is achieved only when two oxoglutarate molecules bind simultaneously, ensuring that full activation occurs only when concentration changes reach a necessary threshold.

In contrast, LTE_4_, a lipid mediator that preferentially remains associated with the membrane lipids, can easily access the alternative site and efficiently activate the receptor, thereby mediating inflammatory responses. This alternative activation occurs with high affinity and efficiency, contributing to the overactivation of the receptor and leading to diseases such as asthma and allergies ^7,14,17,20^. Moreover, the DRY-to-FRY mutation in GPR99/OXGR1 may have evolved to accommodate activation signals from LTE_4_ at the alternative pocket. As the third cysteinyl leukotriene receptor (CysLT_3_R), GPR99/OXGR1 is highly expressed in kidney cells ^1^, fetal brain cells, fibroblast ^33^, as well as immune-related cells including mast cells, eosinophils, and respiratory epithelial mucosa cells ^6,7,9,34^. GPR99/OXGR1 is associated with normal kidney pH-maintaining functions, central nervous system development, fibrosis pathogenesis, allergies and inflammation. Under different physiological contexts, ligand availability and expression levels of GPR99/OXGR1 and G proteins vary, affecting the activation and signaling of the receptor. For instance, in inflammatory microenvironments rich in leukotrienes, the sustained presence of LTE_4_ likely stabilizes its binding to the alternative pocket of GPR99/OXGR1, amplifying Gαi signaling to exacerbate inflammation through leukocyte activation ^7,11,35^. Moreover, an enrichment of GPR99/OXGR1 expression on the respiratory epithelial cells further exacerbates the inflammatory condition. An early study on LTE_4_-induced bronchoconstriction suggested that with a high stability of LTE_4_ compared to LTC_4_ and LTD_4_, asthma develops due to the chronic effect of receptor/ligand-related hyperresponsiveness ^15^. Conversely, in renal tubules, high oxoglutarate concentrations promote Gαq-mediated IP1 accumulation, regulating acid-base homeostasis in kidney ^4^. Acidic renal pH may enhance the polarity of oxoglutarate, thereby favoring Gq signaling through the activation of GPR99/OXGR1 ^4,8^. These context-dependent activation features highlight how physiological environments may shape GPR99/OXGR1 signaling.

LTE_4_ is a high-affinity agonist of GPR99/OXGR1, based on pharmacological characterization of the receptor as a gene product in transfected cell lines ^9^. As a result of these studies, GPR99 is also named CysLT_3_R ^5,9^. How LTE_4_ binds to a GPCR belonging to a different subfamily of the other two CysLT receptors remains unclear. In previous structural studies of antagonist-bound CysLT_1_R and CysLT_2_R, the endogenous agonists CysLTs LTC_4_ and LTD_4_ were proposed to bind to the orthosteric pocket just as the antagonists, which was validated by functional assays ^19,20^. A recent work of the LTD_4_-bound CysLT_2_R structure, published while this study was under revision, provides a solid proof of the binding mode of LTD_4_ to its cognate CysLT_2_R receptor ^21^. LTD_4_ binds within the CysLT_2_R transmembrane pocket, with its polar head reaching to the polar top of the TM pocket and nonpolar interactions spanning TM helices TM4 and TM5 stabilizing its bulky tail (Supplementary Fig. S7a-c). The long bulky tail of LTD_4_ can therefore extend through an orifice opening formed between TM4 and TM5 of CysLT_2_R, similar to the previously proposed binding mode for CysLT_1_R and CysLT_2_R ^26^. In contrast, the GPR99/OXGR1 TM pocket is much smaller, more polar and more hydrophilic, and has no cleft space between TM4 and TM5 for the accommodation of the long bulky tail of LTE_4_, making it poorly suited for LTE_4_ binding (Supplementary Fig. S7d-f). While the superimposed active-state structures of GPR99/OXGR1 and CysLT_2_R are generally similar, the TM3 of CysLT_2_R slightly shifts outwards (Supplementary Fig. S7g), and their binding pockets are very different. By examining the geometry of residues surrounding the TM binding pocket of CysLT_2_R, an opening is observed around the hydrophobic tail of LTD_4_. For the superimposed GPR99/OXGR1 structure at this site, larger amino acid side chains are found to block such an opening (Supplementary Fig. S7h). Despite the ability of being activated by LTE_4_, the sequence of GPR99/OXGR1 is quite different from CysLT_1_R and CysLT_2_R receptors, with less than 30% sequence identity (Supplementary Fig. S7i) ^3,5,9^. Besides, as CysLT_1_R and CysLT_2_R can both selectively recognize LTC_4_ and LTD_4_ with high affinity (but only low affinity for LTE_4_), it is plausible that the binding mode of LTE_4_ is distinct from that of LTC_4_ and LTD_4_ to the respective cysteinyl leukotriene receptors ^36,37^. Differing from Gq coupling of GPR99/OXGR1 when stimulated with oxoglutarate, the receptor is found to activate both Gq and Gi signaling pathways when stimulated with LTE_4_, thus contributing to its proinflammatory and allergy-inducing properties ^7,17^. These results highlight the common and distinct features in the signaling and activation of cysteinyl leukotriene receptors from structural and functional perspectives ^38^. While our studies provide a solid structural foundation, further physiological and pharmacological research is required to advance the development of new cysteinyl leukotriene receptor-targeting drugs.

GPCR drug discovery now marches beyond the canonical transmembrane pocket, with increasing efforts focusing on allosteric ligands that bind to alternative binding sites and modulate receptor functions with signaling bias ^39–44^. Compared with GPR99/OXGR1, several other GPCRs, including S1PR3, GPR88, GPR35, GPR174, PTHR and CB2, share similar alternative allosteric sites and mechanisms. For instance, the sphingosine-1-phosphate (S1P) receptor S1PR3 can be allosterically modulated by ligands binding to its extracellular vestibule (S1P, and its analogs) or deep within the transmembrane pocket (CYM-5541, CBP-307, BAF-312, FTY720-P, and VPC23019), fine-tuning its response and influencing G protein coupling bias towards Gi, Gq and G13 ^45–47^. For an important GPCR regulating neurological functions, GPR88, its synthetic agonist 2-PCCA binds to a cavity at the cytosolic ends of TM5 and TM6 close to ICL3, directly interacts with the α5 helix of Gi and further stabilizes the receptor-Gi complex ^48^. Further exploration of the alternative site on GPR99/OXGR1 may offer a strategy to achieve biased agonism and antagonism, with biased modulators that can selectively bind to different binding pockets and stabilize conformations that favor either Gi or Gq coupling in different physiological environments.

In summary, the present study provides structural insights into oxoglutarate binding to GPR99/OXGR1. The characteristic binding of two oxoglutarate molecules simultaneously may help to address how a GPCR recognizes the small and charged dicarboxylates with high specificity but low efficacy. Despite our effort in resolving the LTE_4_-bound GPR99/OXGR1 structure, no EM density was found in the orthosteric binding pocket as originally expected. Results from a multitude of experiments and comparative analysis indicate that the TM pocket in GPR99/OXGR1 is too small to accommodate the bulky cysteinyl leukotriene. It is likely that LTE_4_ uses an alternative binding site for activation of GPR99/OXGR1, which has been clearly identified as CysLT_3_R through rigorous pharmacological experiments. The present work could not conclusively identify the alternative site for LTE_4_ binding due to poor EM density, but the collected information may help further exploration of the LTE_4_ binding mechanism. Further structural and functional characterization of the alternative binding site on GPR99/OXGR1 will be crucial to unlock its therapeutic potential.

## METHODS

### Design of constructs

The cDNA encoding human wild-type GPR99 was synthesized by General Biol (Chuzhou, China). The constructs were created by cloning the full-length coding sequence into a pFastbac1 vector. To facilitate protein expression and purification, a hemagglutinin (HA) signal peptide, a FLAG tag, a human rhinovirus 3C (HRV 3C) protease cleavage site (LEVLFQGP) and a thermostabilized apocytochrome b562RIL (BRIL) were added to the N terminus, following our previously reported method ^24^. The Gq protein was designed based on the Gi backbone by inserting the C termini of Gq (residues 332 to 359) to DNGαi1 (dominant negative Gαi1 with G203A and A326S mutation) backbone (residues 1 to 326), which provided an additional site for the single-chain antibody variable fragment scFv16 to stabilize the complex. This strategy and the resulting chimera Gq were widely used, as demonstrated in the reported structure determination of the GPCR-Gq complex ^49–51^. Human Gβ1 and Gγ2 cDNAs with N-terminal 6×His tag were respectively cloned into a pFastBac-Dual vector. scFv16 was fused with a GP67 signal peptide at the N-terminal and an 8× His tag at the C-terminal. For the functional assay, the human GPR99 coding sequence was cloned into the pcDNA3.1 vector. Point mutations were introduced using PCR-mediated site-directed mutagenesis.

### Purification of scFv16

Secreted scFv16 was purified from expression media of baculovirus-infected Sf9 insect cell culture using Ni-NTA. The supernatant from 2 L of culture was collected and loaded onto a gravity column packed with Ni-NTA resin. The resin was washed with 20 mM HEPES pH 7.5, 500 mM NaCl, and the protein was eluted with 20 mM HEPES pH 7.5, 150 mM NaCl and 250 mM imidazole. Eluted protein was concentrated and loaded onto Superdex 200 increase 10/300 size exclusion column (GE Healthcare Life Sciences, Sweden). The peak fractions were collected and concentrated, rapidly frozen in liquid nitrogen, and stored at −80 °C.

### Expression and purification of the signaling complex

The bac-to-bac baculovirus expression system and Sf9 insect cells were used to express protein for structure determination. The baculoviruses were prepared according to the manual book (Thermo Fisher Scientific). For protein expression, baculoviruses of GPR99, Gαq, and Gβ1γ2 were co-transfected into Sf9 cells when they reached a density of 2×10^6^ cells/mL. The ratio of baculoviruses for transfection was 1:1:1, and then the proteins were expressed over a 48-hour period. The cell culture was harvested by centrifugation at 2000 x g for 15 minutes and kept frozen at −80 °C.

To purify the complex, the ligand (oxoglutarate or LTE4) was added to induce GPR99-Gq complex formation. Cell pellets were resuspended in buffer containing 25 mM HEPES pH 7.4, 50 mM of NaCl, 5 mM of KCl, 5 mM of CaCl2, 5 mM of MgCl2, 5% glycerol, 25 mU/mL apyrase, 2.5 μg/ml leupeptin, 0.16 mg/ml benzamidine, 100 μM of oxoglutarate or 1 μM of LTE4. Cell membranes were collected by centrifugation after 30 minutes incubation. The cell membranes were then solubilized in 20 mM HEPES pH 7.4, 100 mM NaCl, 0.8% LMNG, 0.1% CHS, 10% glycerol, 2.5 μg/ml leupeptin, 0.16 mg/ml benzamidine, 50 μM of oxoglutarate or 1 μM of LTE4. After 2-hour incubation at 4°C, the supernatant was cleared by centrifugation at 20,000 × g for 30 minutes. A gravity-flow column with anti-FLAG affinity resin (GenScript Biotech) was used to capture the complex, followed by washing with 10 column volumes of wash buffer containing 20 mM of HEPES (pH 7.4), 100 mM of NaCl, 5% glycerol, 2 mM of CaCl2, 2 mM of MgCl2, 0.05% (w/v) LMNG, 0.05% (w/v) GDN (Anatrace), 0.003% (w/v) CHS, 20 μM of oxoglutarate or 0.4 μM of LTE4. The protein complex was eluted with buffer containing 20 mM of HEPES (pH 7.4), 100 mM of NaCl, 2 mM of CaCl2, 2 mM of MgCl2, 0.01% (w/v) LMNG, 0.001% (w/v) GDN (Anatrace), 0.001% (w/v) CHS, 0.2 mg/ml FLAG peptide (Sigma-Aldrich), 20 μM of oxoglutarate or 0.4 μM of LTE4. The eluate was concentrated by an Amicon Ultra-15 Centrifugal Filter Unit (Millipore) and separated on a Superose 6 10/300 size-exclusion chromatography column (Cytiva) pre-equilibrated with 20 mM HEPES (pH 7.4), 100 mM NaCl, 0.0015% LMNG, and 0.0005% GDN, 0.0003% CHS, 20 μM of oxoglutarate or 0.4 μM of LTE4. The relevant peak fractions of the protein complex were collected, analyzed on SDS-PAGE, analyzed on western-blotting, concentrated to approximately 10 mg/mL, and stored at -80°C.

### Cryo-grid preparation and EM data collection

Negative stain electron microscopy was performed on all the samples to confirm homogeneity and complex formation. For cryo-grid preparation, aliquots of 3uL of purified protein complex were applied onto an Ultrafoil 300 mesh R1.2/1.3 holy gold (Au) grid, which had already been glow-discharged by Tergeo-EM plasma cleaner. Afterward, the grids were blotted for 3.5 s with a blot force of 1 in 100% humidity at 4 °C to remove excess samples, and then quickly plunged into liquid ethane using a Vitrobot Mark IV (Thermo Fisher Scientific). The prepared grids were stored in liquid nitrogen.

The grid sample screening and data collection were performed using SerialEM software (Mastronarde, 2005). The final data sets were collected by a Gatan K3 direct electron detector installed on a 300 kV Titan Krios G3 microscope. During data collection, a GIF Quantum energy filter (Gatan, USA) was used to exclude inelastically scattered electrons, with the energy slit width set to 20 eV. Movie stacks were acquired at a nominal magnification of 105,000, resulting in a calibrated pixel size of 0.85 Å. The total exposure time was 2.5 s, fractionated into 50 frames at a dose rate of 20.8 e/pixel/s. The defocus range for dataset collection was set from -1.2 to -2.5 μm.

### Image processing and 3D reconstructions

The cryoSPARC version v4.2.1 ^52^ and Relion version 4.0 ^53^ were used to process the cryo-EM datasets. The pipeline was similar to that reported previously ^24^. Motion correction and dose-weighting were applied to align the movie stacks. After contrast transfer function (CTF) estimation, micrographs were manually inspected, and obvious bad micrographs were discarded. Representative particles were then manually picked to generate initial two-dimensional (2D) templates for auto-picking.

For the oxoglutarate-GPR99-Gq dataset, a total of 2,676,089 particles were template-based picked and subjected to 2D classification. After three rounds of 2D classification, particles were selected from the 2D averages with clear secondary features . Ab initio reconstruction was performed by cryoSPARC to generate initial three-dimensional (3D) templates, followed by rounds of 3D classification. Finally, a dataset containing 167,941 particles was used for homogeneous refinement, non-uniform refinement, and local refinement. The global resolution was 3.16 Å, estimated by the ‘gold standard’ criterion (FSC = 0.143).

For the LTE4-GPR99-Gq dataset, template-based particle picking resulted in a dataset of 2,043,778 particles, which was subjected to three rounds of 2D classification. Good particles with clear secondary features in the 2D averages were selected. Then, after multiple rounds of 3D classification, a final dataset containing 100,083 particles was obtained, resulting in a final map with an estimated global resolution of 2.84 Å.

### Model building and refinement

As no experimental GPR99 structures had been reported previously, we used the predicted structure from the AlphaFold Protein Structure Database (AF-Q96P68-F1) as an initial template for model building. The model building of the G protein heterotrimer and scFv16 was facilitated by our previously reported structure (PDB: 8WOG) as the starting template. The model was docked into the electron microscopy density by UCSF chimera ^54^ and manually rebuilt with Coot ^55^, as well as iterative refinement with Phenix ^56^. Final model validation and statistical analysis were performed using Molprobity ^57^. The graphic structural figures were prepared using UCSF Chimera, ChimeraX and PyMOL.

### IP-one accumulation assay

Human GPR99, both in its wild-type form and in mutants, were expressed in HeLa cells 24 hours before harvesting (Invitrogen, L3000001). Gq downstream signaling, which leads to the accumulation of IP1, was measured using the IP-One Gq HTRF kit (Cisbio). Cells were resuspended in the stimulation buffer (Cisbio) and incubated with varying concentrations of oxoglutarate (A610290, Sangon Biotech, Shanghai, China) or leukotriene E4 (HY-113465, Monmouth Junction, NJ) diluted in the stimulation buffer for 30 minutes at 37°C. IP1 accumulation was then assessed according to the manufacturer’s instructions. Fluorescence intensities were recorded using an Envision 2105 multimode plate reader (PerkinElmer). Intracellular IP1 levels were determined based on the fluorescence signals from the samples and IP1 standards.

### cAMP inhibition assay

Human GPR99 (WT/mutants) were expressed in HeLa cells for 24 hrs (Invitrogen, L3000001), and pretreated with 250 ng/mL PTX or vehicle for 12 h at 37°C. Gi downstream signaling, which results in the inhibition of cAMP accumulation was measured. Cells were harvested in HBSS supplemented with 5 mM HEPES, 0.1% (w/v) BSA, and 0.5 mM 3-isobutyl-1-methylxanthine. The plated cells were subsequently stimulated with varying concentrations of chemerin alongside 2.5 μM forskolin for 30 minutes. Intracellular cAMP levels were quantified using the LANCE Ultra cAMP kit (TRF0263, PerkinElmer) according to the manufacturer’s protocol, with readings taken on an EnVision 2105 multimode plate reader (PerkinElmer).

### NanoBiT-based GPR99-G protein dissociation and Gα-Gβγ protein dissociation assays

HEK293T cells were seeded in 24-well plates and incubated for 24 hours. For GPR99-G protein dissociation assay, cells were co-transfected with plasmids encoding GPR99-SmBiT (333 ng/well) and Gαi-LgBiT (167 ng/well). For Gα-Gβγ protein dissociation, cells were co-transfected with plasmids encoding GPR99 (92 ng/well), Gαi-LgBiT (46 ng/well), Gβ (230 ng/well), and SmBiT-Gγ (230 ng/well). Following a 24-hour incubation at 37°C, the transfected cells were harvested and seeded into white 384-well plates. For the luminescence assay, coelenterazine H (Yeasen Biotechnology, Shanghai, China) was added to a final concentration of 10 μM for 1-hour incubation at room temperature, and baseline luminescence signals were measured for 10 minutes using an Envision 2105 multimode plate reader (PerkinElmer). Ligands were then added, and luminescence detection continued for 1 hour.

### Flow cytometry analysis

HEK293T cells were transfected with expression plasmids encoding FLAG-tagged wild-type (WT) or mutant GPR99 for 24 hours at 37°C. The cells were then collected and washed with HBSS containing 0.5% BSA. After washing, the cells were incubated with a FITC-labeled anti-FLAG antibody (Sigma, Cat #F4049; diluted 1:50 in HBSS buffer) for 30 minutes on ice and washed again with HBSS. Flow cytometry (CytoFLEX, Beckman Coulter) was used to quantify the FITC fluorescence signals on cell surfaces. The fluorescence signals were then analyzed to determine the relative expression levels of GPR99 mutants.

### Molecular Docking Analysis

Flexible docking followed by Binding Free Energy calculations were performed using the BIOVIA Discovery Studio (v2019) to evaluate the differential binding affinity of the ligand LTE4 at orthosteric and allosteric sites. Flexible docking in DS is an induced fit protocol to simulate both protein and ligand flexibility. During flexible docking, there are three distinct phases: (i) protein side-chain conformation generation through CHARMm-based ChiFlex method, (ii) generation of ligand conformations and rigid ligand placement into each protein conformation through LibDock, and (iii) induced fit protein flexibility through ChiRotor-based side-chain reconstruction and final pose refinement through CDOCKER. Protein flexibility is introduced in the first and third phases.

Prior to docking, LTE4 was extracted from the cryo-EM structure and preprocessed using the “Prepare Ligands” and “Minimize Ligands” protocols in DS. This step ensured correction of valency errors, adjustment of protonation states, and geometry optimization under the CHARMm force field. The receptor extracted from the Cryo-EM structure was prepared by adding hydrogen, defining the binding site, and specifying flexible side chains for induced fit modeling. Residues within 5 Å of the bound ligand in the cryo-EM structure were selected as flexible for ChiFlex/ChiRotor processing. A docking region was defined using a spherical selection with a radius of 20 Å centered on the geometric center of LTE4 or α-KG. For each receptor, multiple low-energy conformations were generated by ChiFlex. Ligand conformations were generated using the BEST mode in CatConf. Rigid docking poses were generated using LibDock based on binding-site hotspots, which were clustered and filtered before a final refinement stage using CDOCKER.

After the flexible docking, poses were sorted by -CDOCKER_ENERGY, and the top-ranked poses were selected for binding free energy calculations. The “Calculate Binding Energies” section in Discovery Studio estimates the free energy of binding for a receptor-ligand complex by calculating the free energies of the complex, the receptor, and the ligand. Binding free energies were computed using CHARMm-based molecular mechanics and implicit solvation via the PBSA model. Ligand conformational entropy was included in the estimation by enabling the “Estimate Entropy” and “Ligand Conformational Entropy” options. The reported energy terms include van der Waals, electrostatic, ligand strain, receptor strain, and solvation contributions, providing an overall estimate of binding energy for each pose. The poses with the lowest binding energy were selected as the final poses for further analysis.

### Molecular Dynamics (MD) Simulations and Analysis

Molecular dynamics (MD) simulations started with the atomic coordinates extracted from the LTE4-GPR99 cryo-EM structure resolved in this study. The protonation states of the complex were assigned for a pH of 7.4 using the H++ web server. Next, the CHARMM-GUI membrane builder was used to embed the complex within a POPC (1-palmitoyl-2-oleoyl-sn-glycero-3-phosphocholine) lipid bilayer ^58^. The entire system was then solvated in a periodic water box (TIP3P model) containing 0.1 M NaCl. CHARMM36m force field was applied to model the system. Simulations were run on GPUs using GROMACS (version 2024.2) ^59^. After initial energy minimization and equilibration, three independent production simulations were performed, each lasting 1 μs. The stability of the ligand-receptor complex was monitored throughout the simulations by calculating the root mean square deviation (RMSD). Analysis of these trajectories confirmed that LTE4 consistently adopted a stable conformation in the alternative pocket above ICL2 of GPR99.

### Statistical Analysis

The data analysis was performed using Prism 9.5.0 (GraphPad, San Diego, CA). Dose-response curves for agonist analysis were generated using the log[agonist] vs. response equation (three parameters) within the software. For the IP1 and cAMP assay, data points were expressed as percentages (mean ± SEM) relative to the maximum response level for each sample, based on at least three independent experiments, as indicated in the figure legends. For NanoBiT assays, data points were expressed as fold changes (mean ± SEM) relative to negative control (cells treated without ligands) for each sample, based on at least three independent experiments, as indicated in the figure legends. EC50 values were derived from the dose-response curves. For cell surface expression, data points were shown as percentages (mean ± SEM) of the flow cytometry fluorescence signals of wild-type GPR99. Statistical comparisons were made using one-way analysis of variance (ANOVA). A *p*-value of 0.05 or less was considered statistically significant.

## ACKNOWLEDGMENTS

This work was supported by the National Natural Science Foundation of China (grant number: 82402140), The Medical Scientific Research Foundation of Guangdong Province, China (grant number: B2025178), Special Project for Clinical and Basic Sci & Tech Innovation of Guangdong Medical University (grant number: GDMULCJC2024121 and GDMULCJC2024156), Shenzhen Science and Technology Program Grant No. RCBS20221008093330067 (to A. L.). This work was also supported by grants from the National Key R&D Program of China (Grant 2019YFA0906003); Shenzhen Medical Research Fund SMRF-D2403009 (to R.D.Y.), Yunnan Key Research and Development Program Grant 202402AA310032 (to R.D.Y.), China Postdoctoral Science Foundation 2022M713049 (to A. L.) and start-up funds from Dongguan Songshan Lake Central Hospital (grant number: ky-2025-015, to A. L.).

## AUTHOR CONTRIBUTIONS

A.L. and R.D.Y. conceived, initiated, and designed the project. A.L. and R.D.Y. supervised the study. A.L. and Y. L. performed the experiments, analysis the data and prepare the figures. A.L., Y. L. and R.D.Y. wrote the manuscript.

## DATA AVAILABILITY

The atomic coordinates for the oxoglutarate-GPR99-Gq complex and the LTE_4_-GPR99-Gq complex have been deposited in the Protein Data Bank with accession codes 8YYW and 8YYX, respectively. The corresponding EM maps have been deposited in the Electron Microscopy Data Bank with accession codes EMD-39681 and EMD-39682, respectively. All data needed to evaluate the conclusions in the paper are present in the main text or the supplementary materials.

