## Supplementary Information for "Molecular insights into ligand recognition and signaling of GPR99/OXGR1"

Supplementary Fig S1-S12

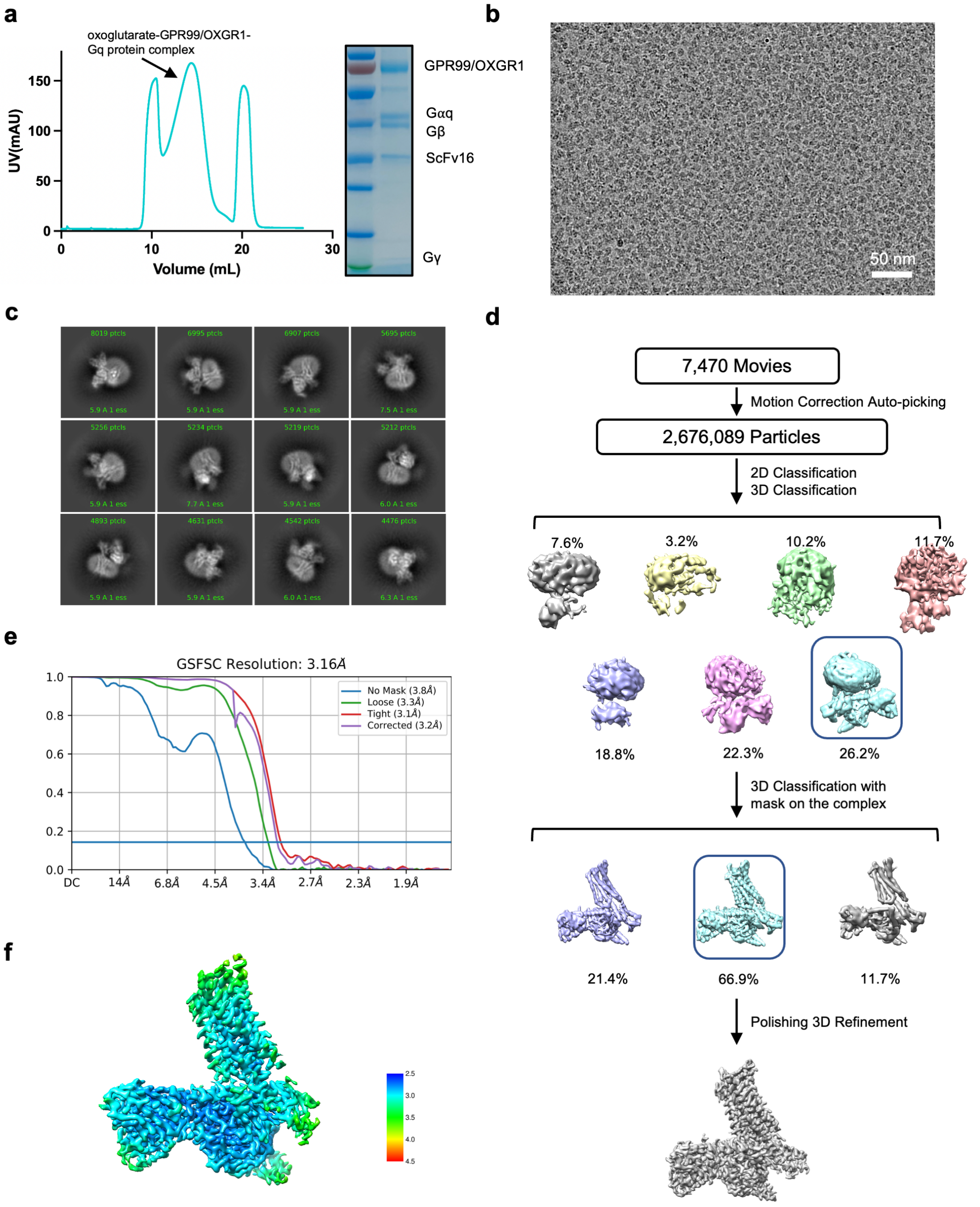


**Supplementary Fig. S1 Purification and cryo-EM data processing of the oxoglutarate-bound GPR99/OXGR1-Gq complex. a,** Size-exclusion chromatography profile and SDS-PAGE analysis of the GPR99/OXGR1-Gq complex bound with oxoglutarate. **b,** Representative micrograph after motion correction and dose weighting. **c,** 2D class averages of the GPR99/OXGR1-Gq complex bound with oxoglutarate. **d,** Workflow of cryo-EM data processing using cryoSPARC. **e,** Gold standard Fourier shell correlation (FSC) curve, indicating an overall nominal resolution of 3.16 Å. **f,** Local resolution map.

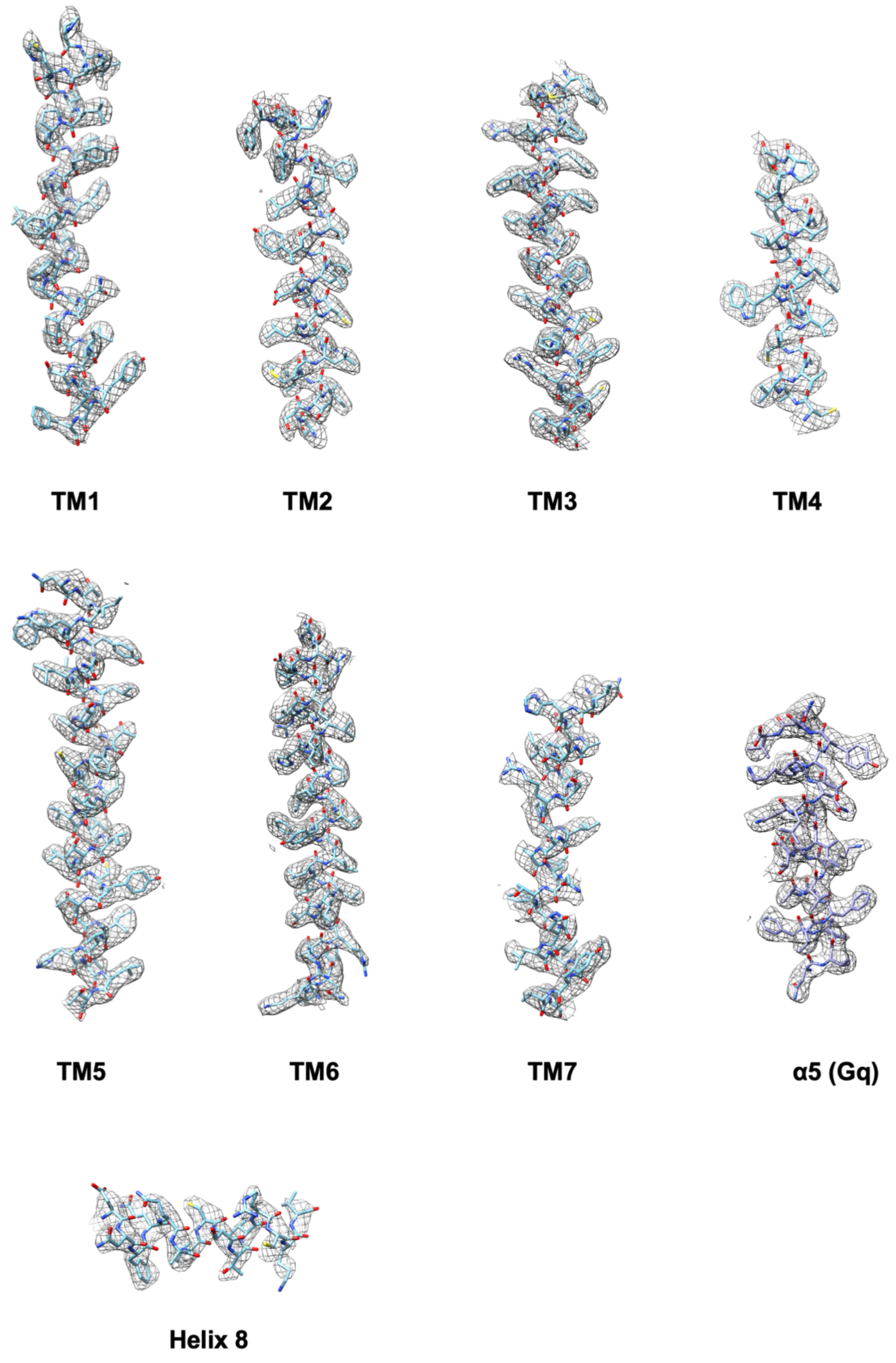


**Supplementary Fig. S2 Representative density maps and models of the oxoglutarate-bound GPR99/OXGR1-Gq complex.**

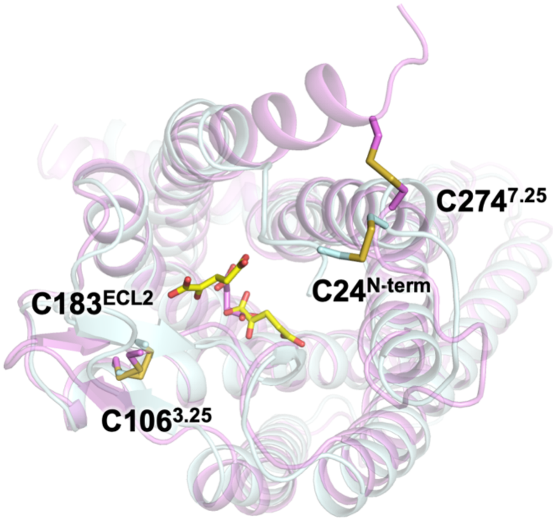


**Supplementary Fig. S3 Conserved disulfide bridges in GPR99/OXGR1 (cyan) and GPR91/SUCNR1 (magenta, PDB ID: 8WOG).** Disulfide bonds are highlighted in yellow, with the corresponding cysteine residues labeled on GPR99/OXGR1. The oxoglutarate ligands for GPR99/OXGR1 are shown as yellow sticks, and the succinate ligand for GPR91/SUCNR1 is shown as pink sticks.

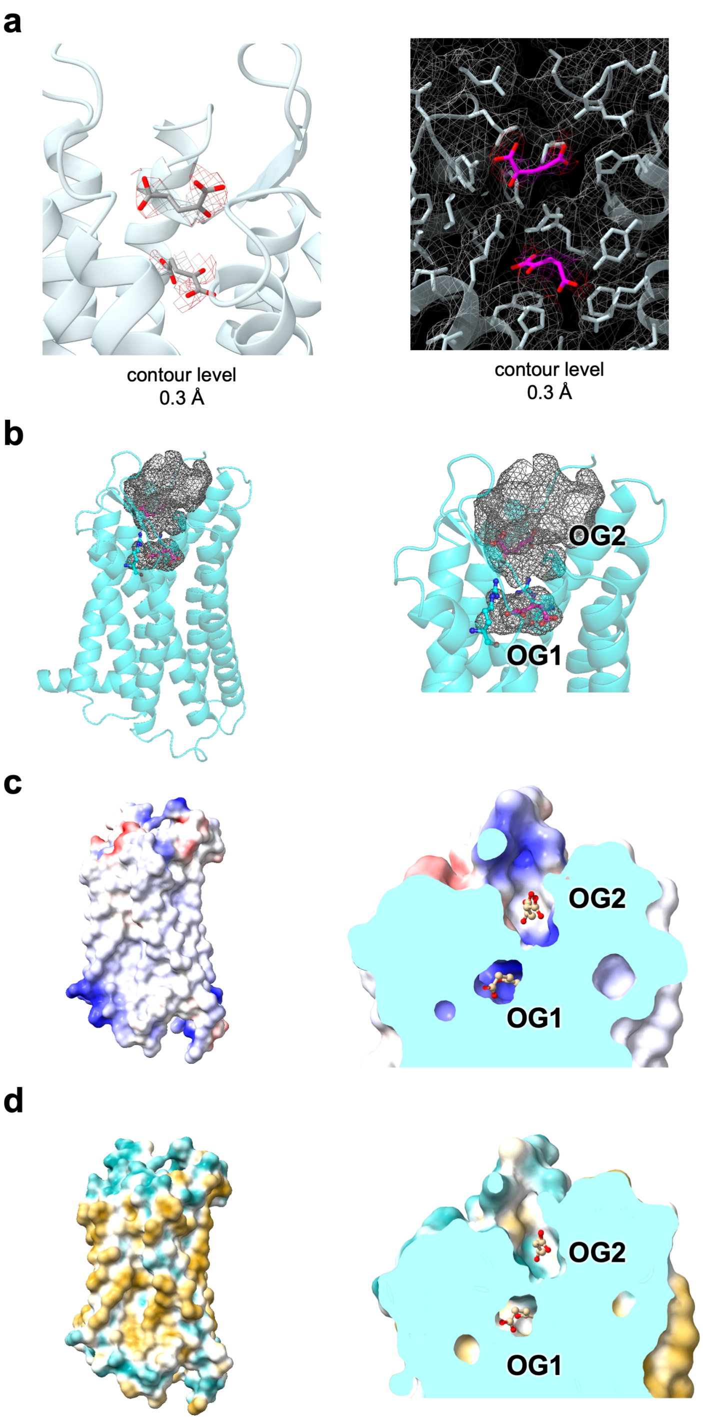


**Supplementary Fig. S4 The transmembrane binding pocket for oxoglutarate in GPR99/OXGR1. a,** EM map density of oxoglutarate molecules at a contour level of 0.3 Å. Left, the EM map density of the two oxoglutarate molecules. Right, the EM map density of the oxoglutarate molecules (colored in magenta) and the GPR99 TM binding pocket (colored in light cyan) in a different angle. **b,** The cavity in the transmembrane binding pocket of GPR99/OXGR1. The cavity is shown as meshed grey surface, and GPR99/OXGR1 is represented as cyan cartoon. **c,** Electrostatic potential surface of the GPR99/OXGR1 binding pocket for oxoglutarate. Red indicates negative charges and blue stands for positive charges. Oxoglutarate molecules are shown in sticks. **d,** Hydrophobicity surface of the GPR99/OXGR1 binding pocket for oxoglutarate. Yellow represents hydrophobic regions and cyan indicates hydrophilic regions.

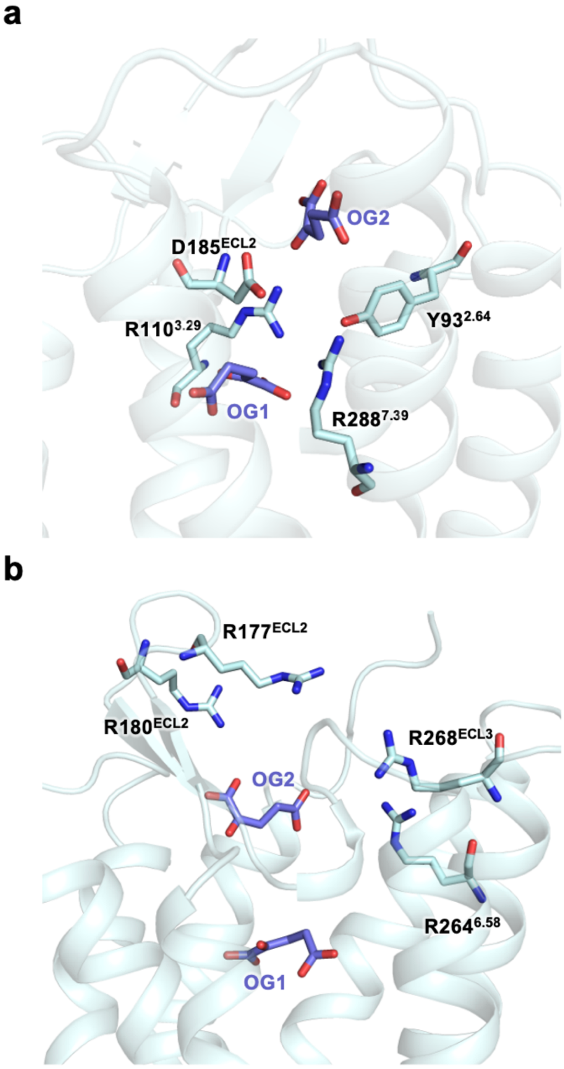


**Supplementary Fig. S5 Arginine residue network gating the transmembrane binding pocket of GPR99/OXGR1. a,** Separation of OG1 and OG2 binding cavity by R110$^{3.29}$, R288$^{7.39}$, Y93$^{2.64}$ and D185$^{ECL2}$. **b,** Gating of the transmembrane binding pocket of GPR99/OXGR1 by arginine residues on ECL2 and ECL3. OG1 and OG2 are marked as purple sticks.

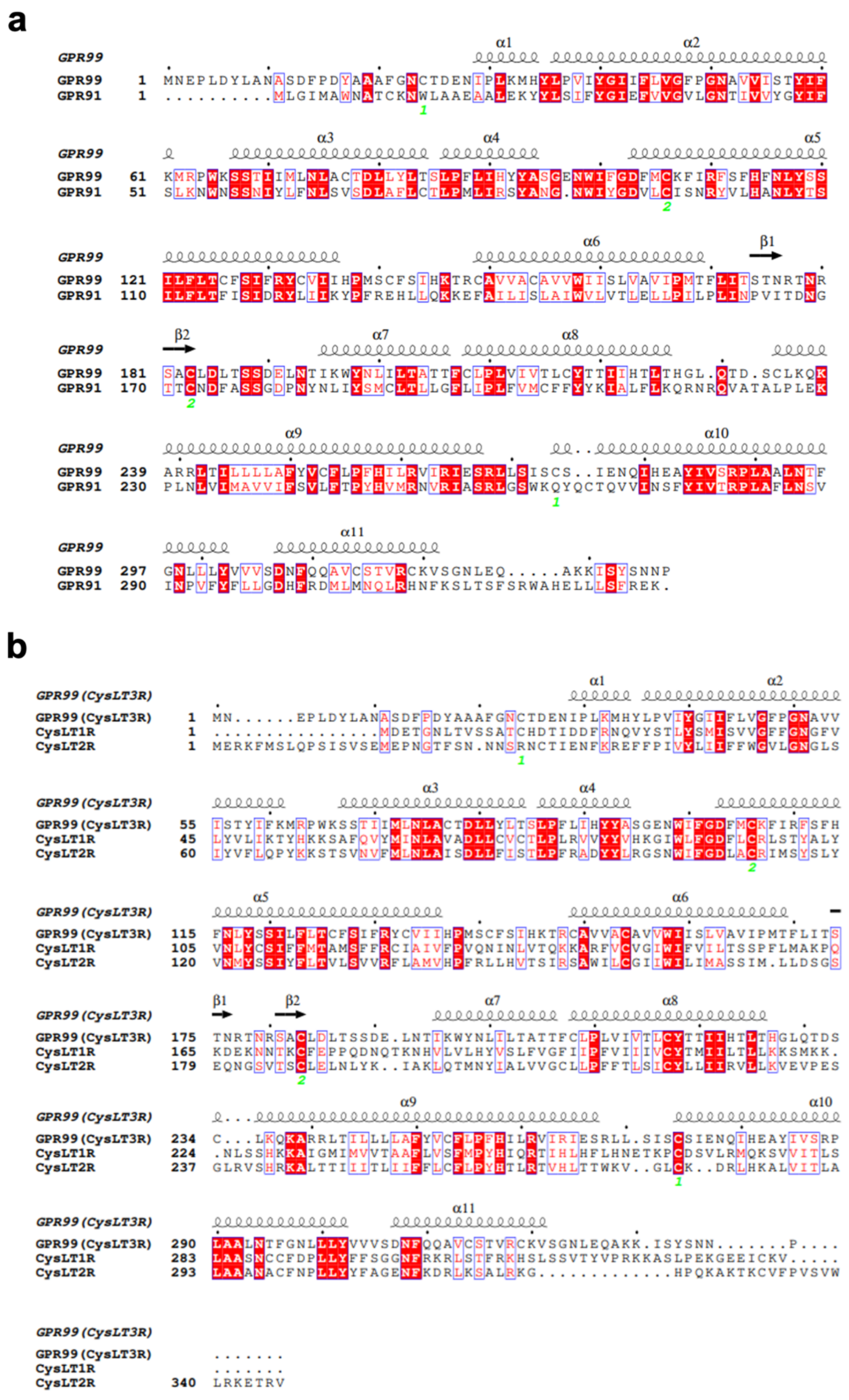


**Supplementary Fig. S6 Full sequence alignment of dicarboxylate receptors and cysteinyl leukotriene receptors. a,** Sequence alignment of human oxoglutarate receptor GPR99/OXGR1 and human succinate receptor GPR91. **b,** Sequence alignment of human cysteinyl leukotriene receptor GPR99/OXGR1 ($CysLT_3R$), $CysLT_1R$, and $CysLT_2R$.

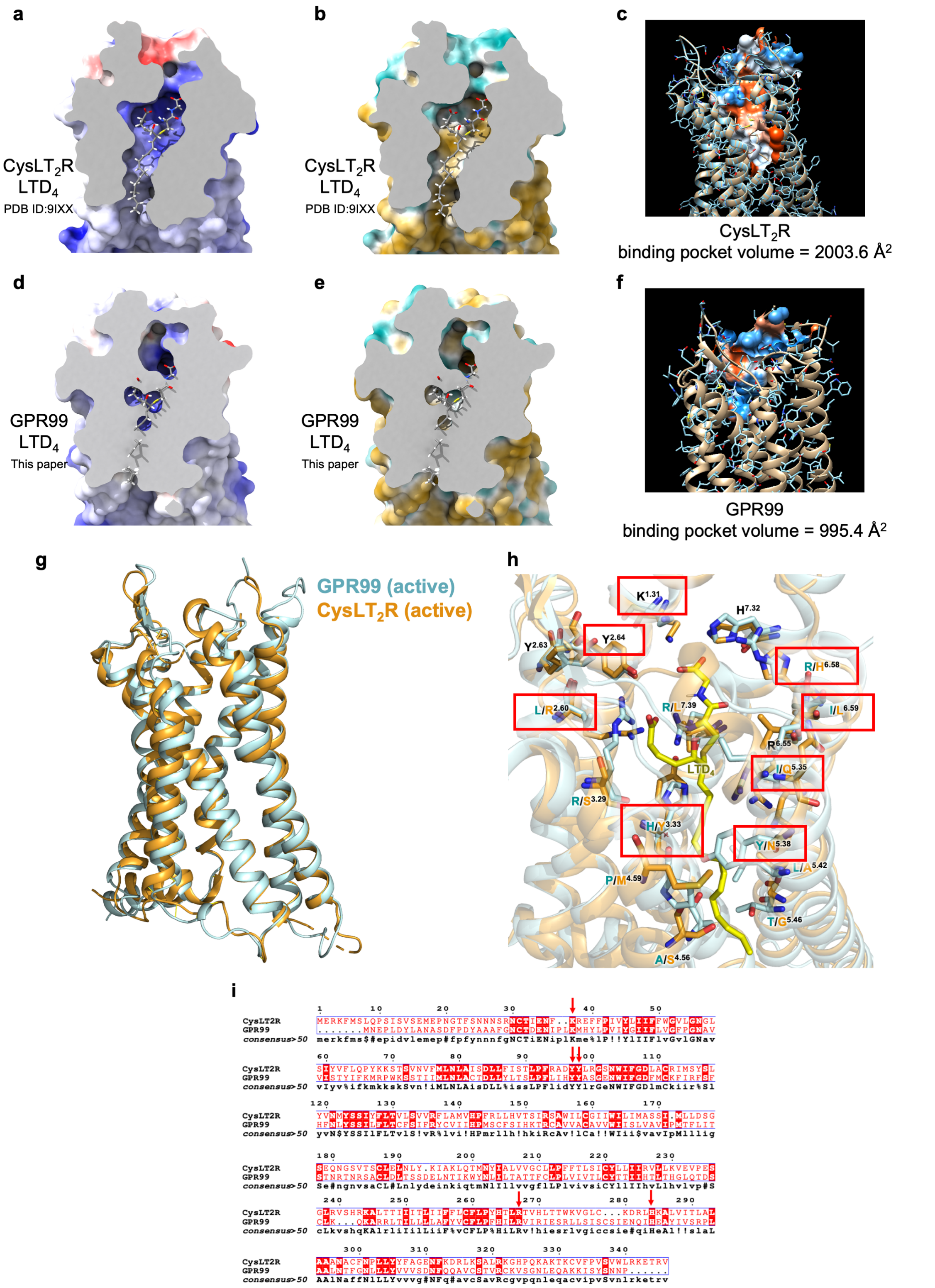


**Supplementary Fig. S7 Comparison of the structures of GPR99/OXGR1 and CysLT$_2$R.** The LTD$_4$-CysLT$_2$R (PDB ID: 9IXX) in an active conformation is used as a reference model. Panels **a, b, d, e** are the receptor structures sliced at the TM binding pocket in the same plane. In panels **d-e**, the $LTD_4$ molecule in the 9IXX model is retained and shown in the aligned receptor structure of GPR99/OXGR1. Panels **a** and **d** show the electrostatic surface of the receptors, and panels **b** and **e** show the hydrophobicity surface of the receptors. In panels **c** and **f**, the volumes of TM binding pocket are measured using CASTpFold. The computed binding pockets are shown as surface. **g,** overall superimposed structures of GPR99/OXGR1 (cyan, this paper) and $CysLT_2R$, both in active state. For display clarity, the ligands are removed from the orthosteric pocket. **h,** the transmembrane (TM) binding pocket of $CysLT_2R$ with bound $LTD_4$ (yellow) and the superimposed structure of GPR99/OXGR1. Notably, $CysLT_2R$ residues important for interaction with $LTD_4$ are either not present in GPR99/OXGR1 or changed to amino acids of different properties. For example, L188, E189 and L190 in the ECL2 of $CysLT_2R$, that interact with $LTD_4$, are not found in the equivalent position in GPR99/OXGR1. Critical amino acids of $CysLT_2R$ that form hydrogen bonds or hydrophobic interaction with $LTD_4$ are either shifted in position (K1.31, Y2.64, H7.32) or substituted with different amino acids (R2.60 to L, Y3.33 to H, /Q5.35 to I, N5.38 to Y, H6.58 to R, and L7.39 to R) in GPR99/OXGR1. **i,** sequence alignment of GPR99/OXGR1 and $CysLT_2R$. Conserved sites in the TM binding pockets are indicated with a red arrow.

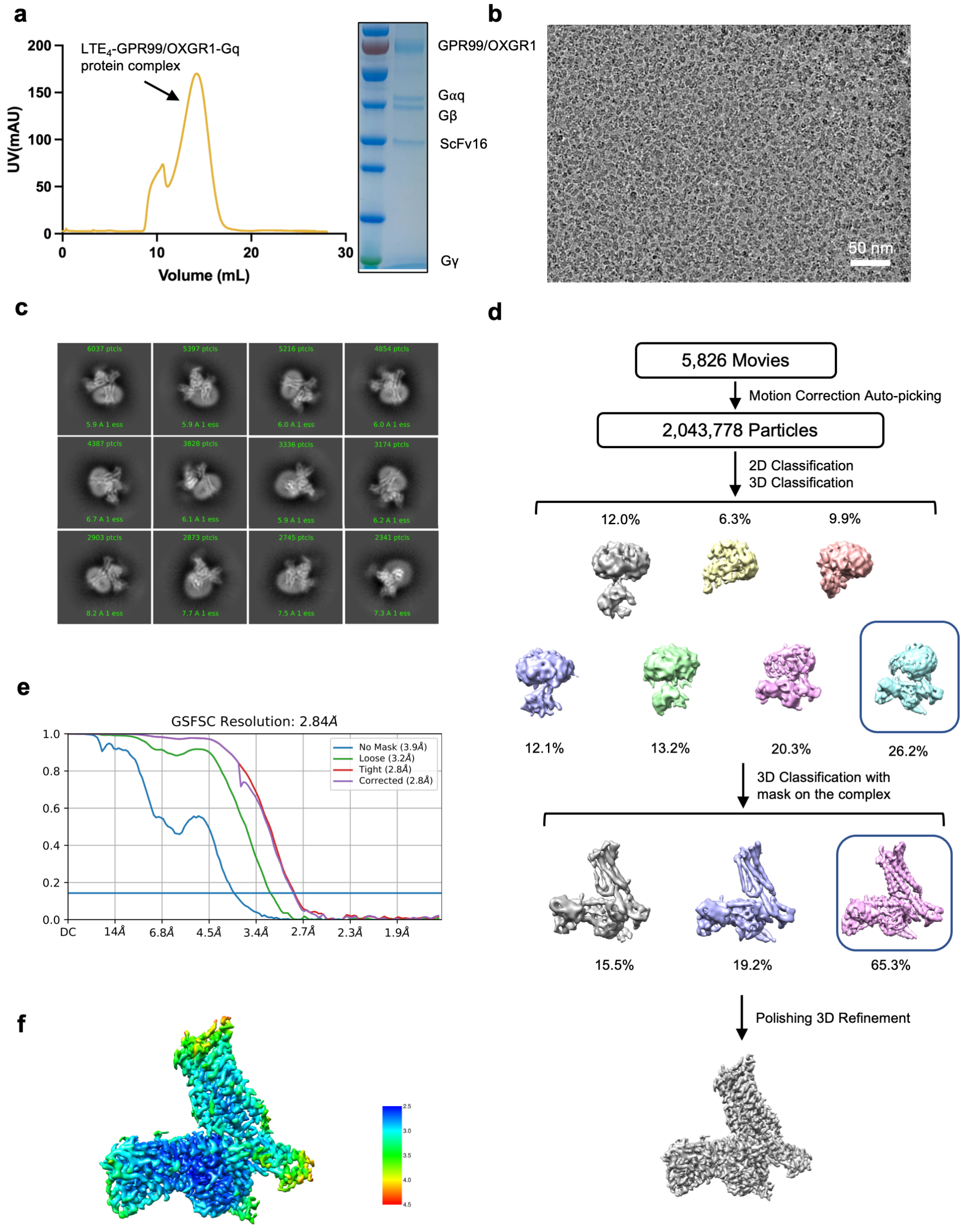


**Supplementary Fig. S8 Purification and cryo-EM data processing of the LTE4-bound GPR99/OXGR1-Gq complex. a,** Size-exclusion chromatography profile and SDS-PAGE analysis of the GPR99/OXGR1-Gq complex bound with LTE4. **b,** Representative micrograph after motion correction and dose weighting. **c,** 2D class averages of the GPR99/OXGR1-Gq complex bound with LTE4. **d,** Workflow of cryo-EM data processing using cryoSPARC. **e,** Gold standard Fourier shell correlation

(FSC) curve indicates an overall nominal resolution of 2.84 Å. **f,** Local resolution map.

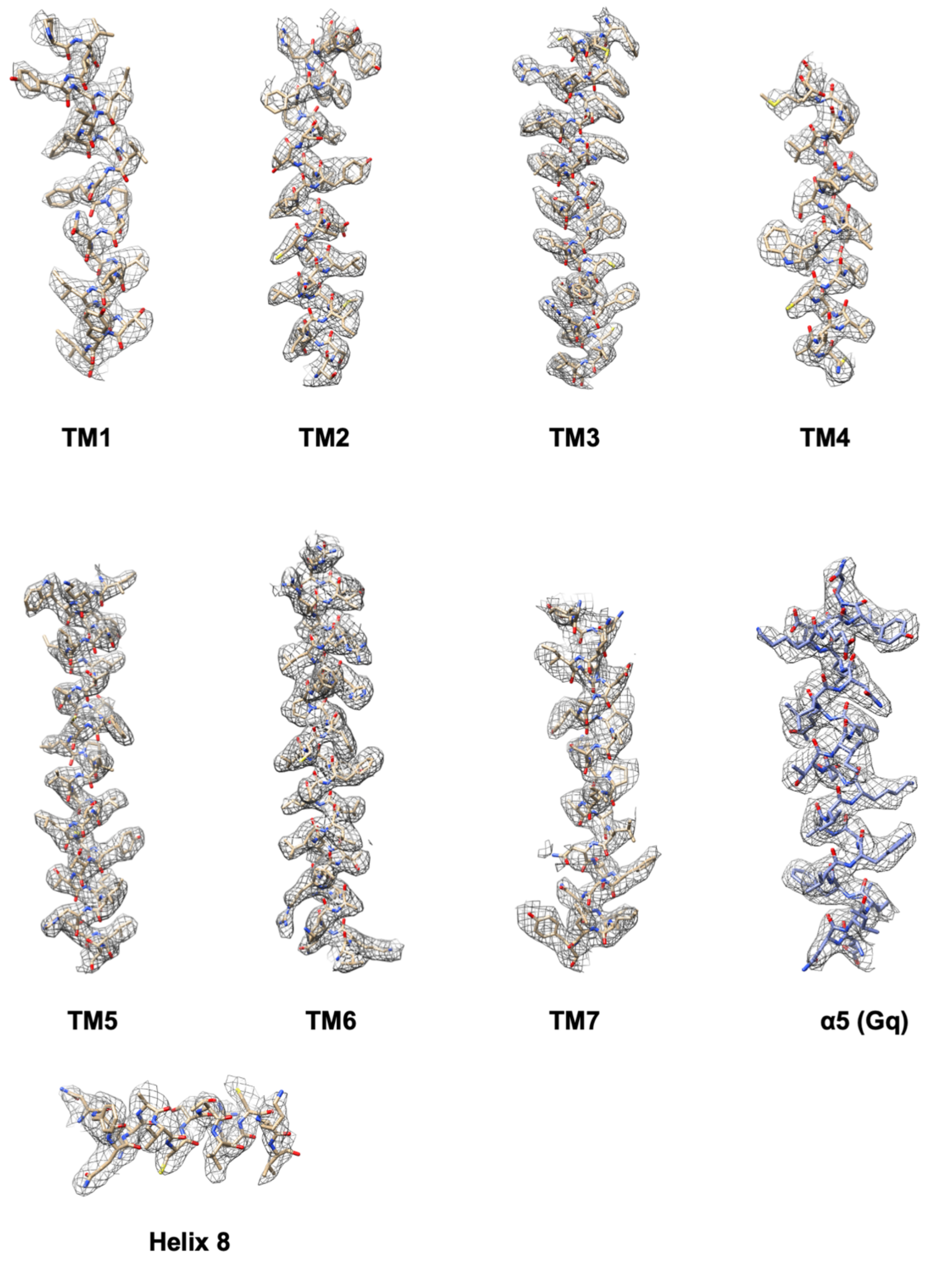


**Supplementary Fig. S9 Representative density maps and models for the $LTE_4$-GPR99/OXGR1-Gq complex.**

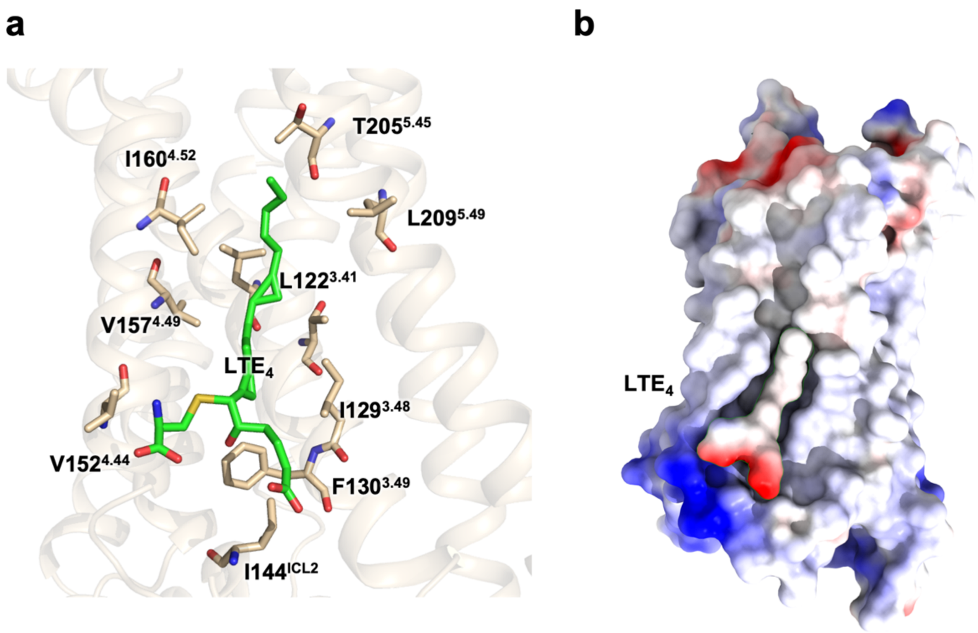


**Supplementary Fig. S10 The non-canonical binding pocket of $LTE_4$ in GPR99/OXGR1. a,** Hydrophobic interactions between $LTE_4$ and GPR99/OXGR1. Hydrophobic residues are shown as champagne-colored sticks. **b,** Electrostatic potential surface of $LTE_4$-GPR99/OXGR1 interaction. The negatively charged head of $LTE_4$ (red) docks into the positively charged ICL2 cavity.

**a**

| CDOCKER binding energy (kcal/mol) | GPR99/OXGR1 TM pocket | GPR99/OXGR1 alternative binding site |
|---|---|---|
| 1 | -7.4703 | -37.7072 |
| 2 | -6.244 | -36.7584 |
| 3 | -4.6715 | -36.5264 |

**b**

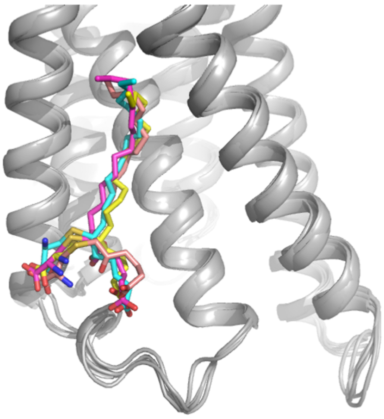

**c**

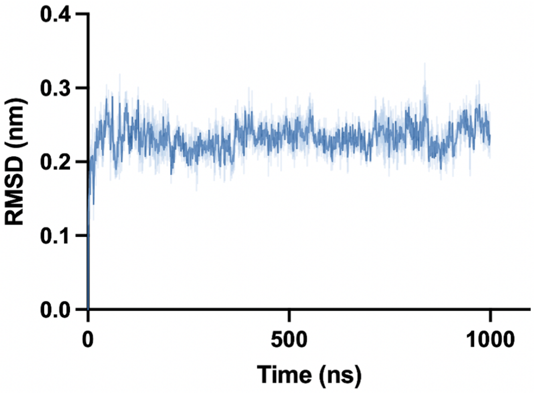


**Supplementary Fig. S11 Molecular docking and MD simulations of $LTE_4$ binding to GPR99/OXGR1. a,** molecular docking-computed binding energies of $LTE_4$ docked to the orthosteric binding site and the alternative binding site above ICL2 of GPR99/OXGR1, respectively. Three independent binding poses were generated for binding energy computation. **b,** stable poses of $LTE_4$ in the alternative binding site from 3 independent 1-μs MD simulations (colored in magenta, salmon pink and yellow), superimposed with $LTE_4$ (cyan) in the $LTE_4$-GPR99/OXGR1-Gq structure. **c,** root mean square distance (RMSD) of $LTE_4$ from three independent 1-μs MD simulations.

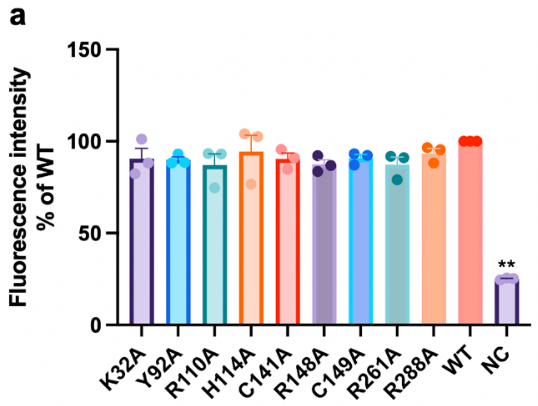


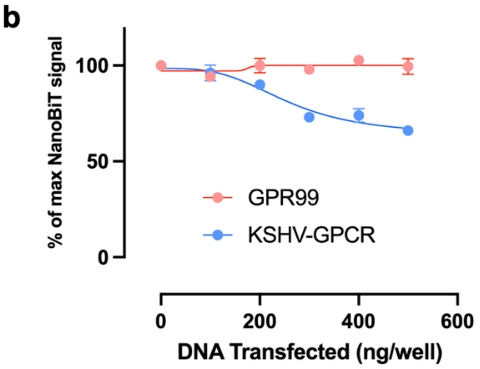


**Supplementary Fig. S12 Expression levels of GPR99/OXGR1 mutants and NanoBiT-based basal activity analysis of GPR99/OXGR1. a**, HEK293T cells were transfected with the plasmids for WT or mutant GPR99/OXGR1 carrying an N-terminal FLAG tag. Flow cytometry analysis were performed using an anti-FLAG M2 antibody. Fluorescence intensities were calculated, and shown as % of the WT GPR99. Data in triplicates were obtained from three independent experiments and plotted. **b**, The basal activity of GPR99/OXGR1 was compared with a GPCR known for constitutive activation (KSHV-GPCR) as a function of increasing receptor expression levels (more DNA used in cell transfection). The NanoBiT-based G protein dissociation assay was used and dissociation of the G protein subunits (α from βγ, with reduced values) was an indicator of receptor activation. Data shown were values from five independent experiments, each with triplicates.
